# Structural insights into TRPV4-Rho GTPase signaling complex function and disease

**DOI:** 10.1101/2023.03.15.532784

**Authors:** Do Hoon Kwon, Feng Zhang, Brett A. McCray, Meha Kumar, Jeremy M. Sullivan, Charlotte J. Sumner, Seok-Yong Lee

**Author notes:** Correspondence to: S.-Y. Lee. These authors contributed equally (Do Hoon Kwon, Feng Zhang).

## Abstract

Crosstalk between ion channels and small GTPases is critical during homeostasis and disease^1^, but little is known about the structural underpinnings of these interactions. TRPV4 is a polymodal, calcium-permeable cation channel that has emerged as a potential therapeutic target in multiple conditions^2–5^. Gain-of-function mutations also cause hereditary neuromuscular disease^6–11^. Here, we present cryo-EM structures of human TRPV4 in complex with RhoA in the apo, antagonist-bound closed, and agonist-bound open states. These structures reveal the mechanism of ligand-dependent TRPV4 gating. Channel activation is associated with rigid-body rotation of the intracellular ankyrin repeat domain, but state-dependent interaction with membrane-anchored RhoA constrains this movement. Notably, many residues at the TRPV4-RhoA interface are mutated in disease and perturbing this interface by introducing mutations into either TRPV4 or RhoA increases TRPV4 channel activity. Together, these results suggest that the interaction strength between TRPV4 and RhoA tunes TRPV4-mediated calcium homeostasis and actin remodeling, and that disruption of TRPV4-RhoA interactions leads to TRPV4-related neuromuscular disease, findings that will guide TRPV4 therapeutics development.

## Main

Although typically considered part of distinct signaling pathways, functional interactions between ion channels and small GTPases have been reported for over two decades and control a range of fundamental processes including cell migration, tumor vascularization, smooth muscle contraction, and mechanosensation, among others^1,12–16^. Insight into how this dynamic crosstalk is achieved is limited by the absence of resolved channel-GTPase structures and structure-guided functional studies.

TRPV4, expressed in the plasma membrane of a wide range of cell types, is a polymodal ion channel whose gating is controlled by multiple endogenous and exogenous stimuli including synthetic ligands, cell swelling, shear stress, and moderate heat^17–19^. TRPV4 mediates calcium dependent regulation of vascular tone, osmoregulation, bone homeostasis, itch, adipose thermogenesis, inflammation, and pulmonary and renal functions^2–4,20^. TRPV4 channel activation is linked to numerous disease states: pulmonary edema and cancer metastasis, amongst other diseases^5,21^. Furthermore, gain-of-function missense mutations cause TRPV4 channelopathies, which are grouped into autosomal dominant neuromuscular disorders (Charcot-Marie-Tooth disease type 2C and distal spinal muscular atrophies) and skeletal disorders (skeletal dysplasias and osteoarthropathy)^6–8,11^. Notably, while skeletal dysplasia mutations are distributed throughout the TRPV4 channel, neuromuscular disease-causing mutations (referred to hereafter as neuropathy mutations) are primarily localized to a confined region of the N-terminal cytoplasmic domain. We previously showed that the cytoskeleton remodeling small GTPase RhoA functionally interacts with TRPV4^22^, but this interaction appears to be perturbed by neuropathy-causing mutations resulting in increased TRPV4 channel activity, cytoskeletal remodeling, and cell process retraction^22^. Overexpression of RhoA suppresses wild type (WT) TRPV4 channel-mediated calcium influx in cultured mouse motor neuron–neuroblastoma fusion (MN-1) cells in response to hypotonicity demonstrating its ability to modulate TRPV4 function (Fig. 1a,b) ^22^.

**Fig. 1.**
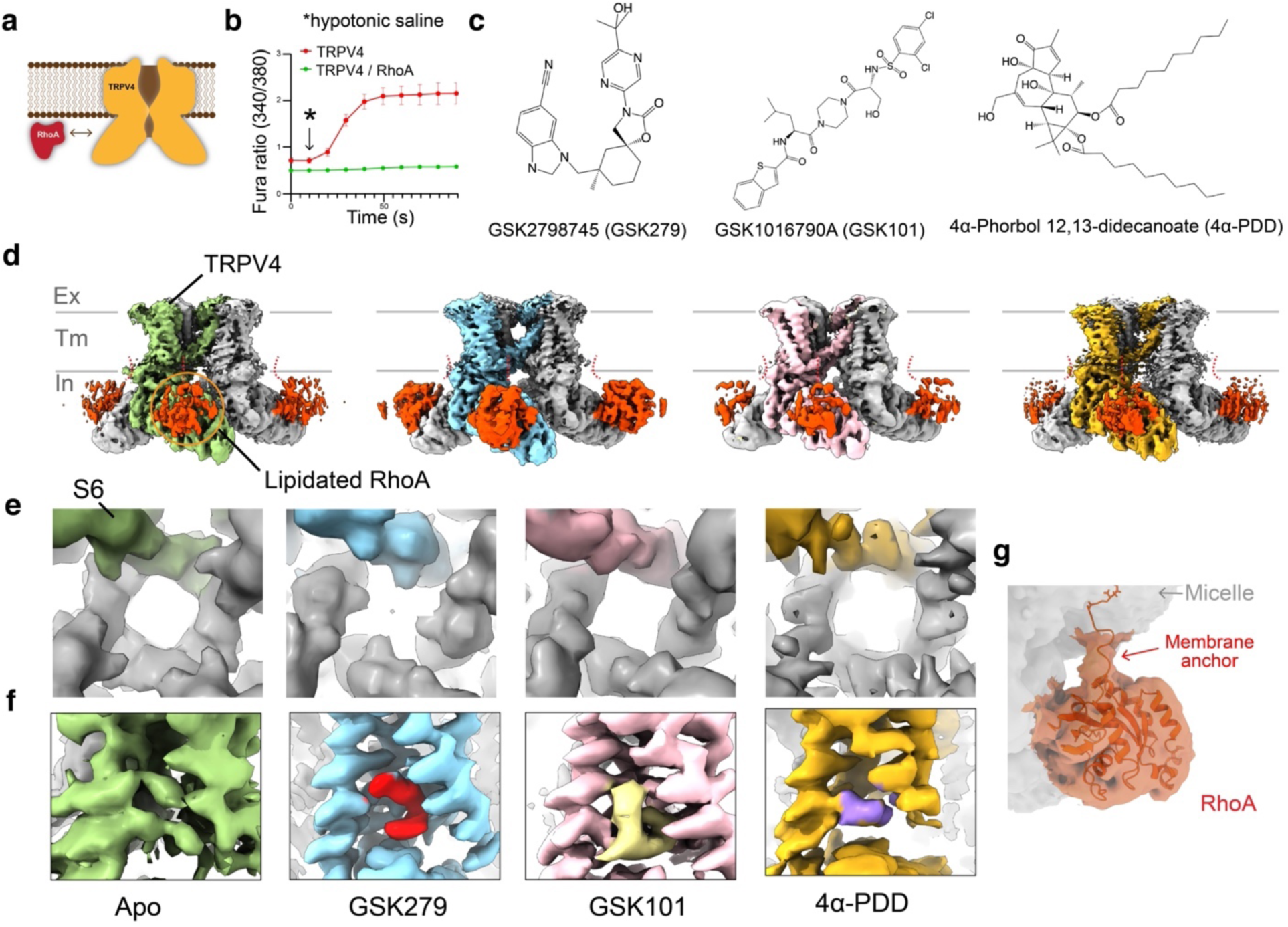
Functional characterization and structure determination of the human TRPV4 – RhoA complex. **a**, Schematic drawing of the functional TRPV4-RhoA interaction. **b**, Averaged calcium imaging traces before and after hypotonic stimulation, denoted by the arrow. Expression of WT TRPV4 alone causes elevated baseline and stimulated calcium influx relative to the co-expression of TRPV4 WT and RhoA. Data are presented as means ± SEM. **c**, Chemical structures of GSK1016790A (GSK101), GSK2798745 (GSK279), and 4α-Phorbol 12,13-didecanoate (4α-PDD). **d**, Cryo-EM structures of hTRPV4-RhoA complex in the apo, closed, open, and 4α-PDD-open states, as indicated. Thresholding, 0.24 (green), 0.25 (cyan), 0.25 (pink), 0.26 (gold) **e**, Close-up view at the S6 gate of 3D reconstructions from (**d**) viewed from the intracellular side, at thresholding 0.14, 0.25, 0.26, and 0.185, respectively. **f**, Close-up view at the ligand binding site of 3D reconstructions from (**d**) at thresholding 0.23, 0.36, 0.34, and 0.22, respectively. **g**, Cryo-EM density of RhoA and lipidated tail of GSK279-hTRPV4-RhoA, viewed from the intracellular side, at thresholding 0.09.

TRPV4 channel inhibition is a promising therapeutic strategy for multiple diseases and conditions^20,23^. Administration of TRPV4 antagonists improves outcomes in animal models of pulmonary edema, blood-retinal and blood-brain barrier breakdown, and peripheral neuropathy^24–26^ and the orally bioavailable TRPV4 antagonist GSK2798745 (Fig. 1c) has been through clinical trials for pulmonary edema, chronic cough, and diabetic macular edema (NCT02119260, NCT02497937, NCT03372603, and NCT04292912)^23,24^. Understanding the structural bases of ligand-dependent TRPV4 gating as well as channel modulation by RhoA will enhance drug design for TRPV4-dependent diseases. To date, the only published structure of TRPV4 is from *Xenopus tropicalis* in its ligand-free, non-conducting state^27^, but the atypical non-domain-swapped tetrameric channel arrangement calls into question its physiological relevance^28^. Therefore, little is known about the agonist- and antagonist-binding sites within TRPV4 and how they exert effects on channel gating.

## Results

### Structures of human TRPV4 in complex with RhoA

We expressed full-length WT human TRPV4 in HEK293S GnTi^-^ cells. The protein was extracted and purified in detergent and the structure determined using single-particle cryo-electron microscopy (cryo-EM) (Figs. 1d-f). Surprisingly, although we overexpressed TRPV4 only, an additional protein density associated with each cytoplasmic domain of the tetrameric TRPV4 channel was resolved in the final three-dimensional (3D) reconstruction (Fig. 1d). The published RhoA crystal structure (PDB: 1FTN) fits reasonably well into this density, suggesting that endogenous RhoA was copurified with overexpressed TRPV4. The presence of RhoA in the final purified TRPV4 samples was confirmed by Western blot using a RhoA-specific antibody (Extended Data Fig. 1a). The C1-symmetric 3D reconstruction showed RhoA occupancy at all four subunits of the TRPV4 homotetramer (Extended Data Fig. 1b). We define this ligand-free TRPV4-RhoA complex structure as the apo state. Three additional complex structures were determined in the presence of distinct ligands (Figs. 1c-f): the antagonist GSK2798745-bound TRPV4-RhoA complex in the closed state, the agonist GSK1016790A-bound TRPV4-RhoA in the open state, and the agonist 4α-phorbol 12,13-didecanoate-bound TRPV4-RhoA in the putative open state (ligands are abbreviated as GSK279, GSK101, and 4α–PDD, respectively). The 3D reconstructions of the four structures were resolved to 3.30 to 3.75 Å resolutions (Extended Data Figs. 1c-h). Particle subtraction followed by focused 3D classification of the transmembrane region and RhoA-bound cytoplasmic domain were performed in turn, resulting in improved map qualities for these regions for model building (Extended Data Fig. 2). Although the EM density for the pore domain in the apo state and the 4α–PDD-bound structures was resolved sub-optimally, the high-quality reconstructions for the GSK279-bound closed state and the GSK101-bound open state enabled us to unambiguously model the register and assign the gate residues (Extended Data Fig. 3). Notably, we observed EM density of the RhoA C-terminus extending to the detergent micelle that surrounds the TRPV4 transmembrane region, suggesting that the geranylgeranylated C-terminus of RhoA anchors to the inner leaflet of the membrane bilayer and facilitates its association with TRPV4 in the cellular context (Fig. 1g).

Human TRPV4 adopts a two-layered homotetrameric architecture where RhoA attaches to the bottom layer. The top layer, or the transmembrane region, comprises the voltage-sensor-like domain (VSLD) and the pore domain. The VSLD is formed by transmembrane helices S1 to S4, while the pore domain contains the S4-S5 junction, S5, the pore helix (PH), the selectivity filter (SF), the pore loop, the pore-lining helix S6, and the TRP domain (Extended Data Figs. 4a,b). The bottom layer is composed of the cytosolic N-terminal ankyrin repeat domain (ARD; comprising six ankyrin repeats, ARs) and a coupling domain (CD), including a helix-turn-helix motif (HTH_CD_) and β-sheets (β_CD_), and a C-terminal domain (CTD) (Extended Data Fig. 4a). RhoA acts as an auxiliary subunit, interacting with the ARD through three loops connecting AR2-AR3, AR3-AR4, AR4-AR5, and with the membrane bilayer through the prenylated C-terminal tail. The apparent stoichiometry of RhoA and each TRPV4 subunit is 1:1 based on the C1 reconstruction (Extended Data Fig. 1b). The human tetrameric TRPV4-RhoA signaling complex structure exhibits a canonical domain-swap tetrameric arrangement where the VSLD of TRPV4 from one subunit interacts with the pore domain from the neighboring subunit (Extended Data Figs. 4b-f). The fold and quaternary structure of full-length human TRPV4 are in stark contrast with the published cryo-EM structure of truncated frog TRPV4, whose pore domains are not swapped (Extended Data Fig. 4g)^27^. Because the domain-swapped architecture is the well-accepted architecture of the TRP channel superfamily^28^, our TRPV4-RhoA structures represent physiologically relevant conformations.

### Agonist- and antagonist-dependent TRPV4 gating

We identified strong and unambiguous EM densities corresponding to GSK279 (antagonist), GSK101 (agonist), and 4α-PDD (agonist) within the cavity between the VSLD and the TRP domain (termed here the VSLD cavity; Figs. 1f, 2a). These compounds are stabilized within the VSLD cavity by many aromatic and polar residues. Due to their higher TMD map quality, we only included the GSK279-bound closed state and the GSK101-bound open state for subsequent analyses of ligand binding and gating.

As the shared binding region for GSK279 and GSK101 complicates functional studies of TRPV4, we utilized the double mutant N456H/W737R (referred to as TRPV4^DM^)^29^ that can be activated by 2-aminoethoxydiphenyl borate (2-APB) (Figs. 2b-e). The putative 2-APB binding site in TRPV4 is located at a distance from the VSLD cavity^30^, thus 2-APB binding to TRPV4^DM^ does not interfere with either GSK279 or GSK101 binding. Y553A, D743A, and F524A mutations on the background of TRPV4^DM^ suppress TRPV4 activation by GSK101 relative to that by 2-APB (Fig. 2c), while D743A and D546A attenuate channel inhibition by GSK279 (Fig. 2e). Previous studies have suggested that the S2-S3 loop, S3, and S4 are involved in TRPV4 sensitivity to 4α– PDD and the inhibitor, HC067047^31,32^, which is consistent with our findings (Extended Data Fig. 5a).

**Fig. 2.**
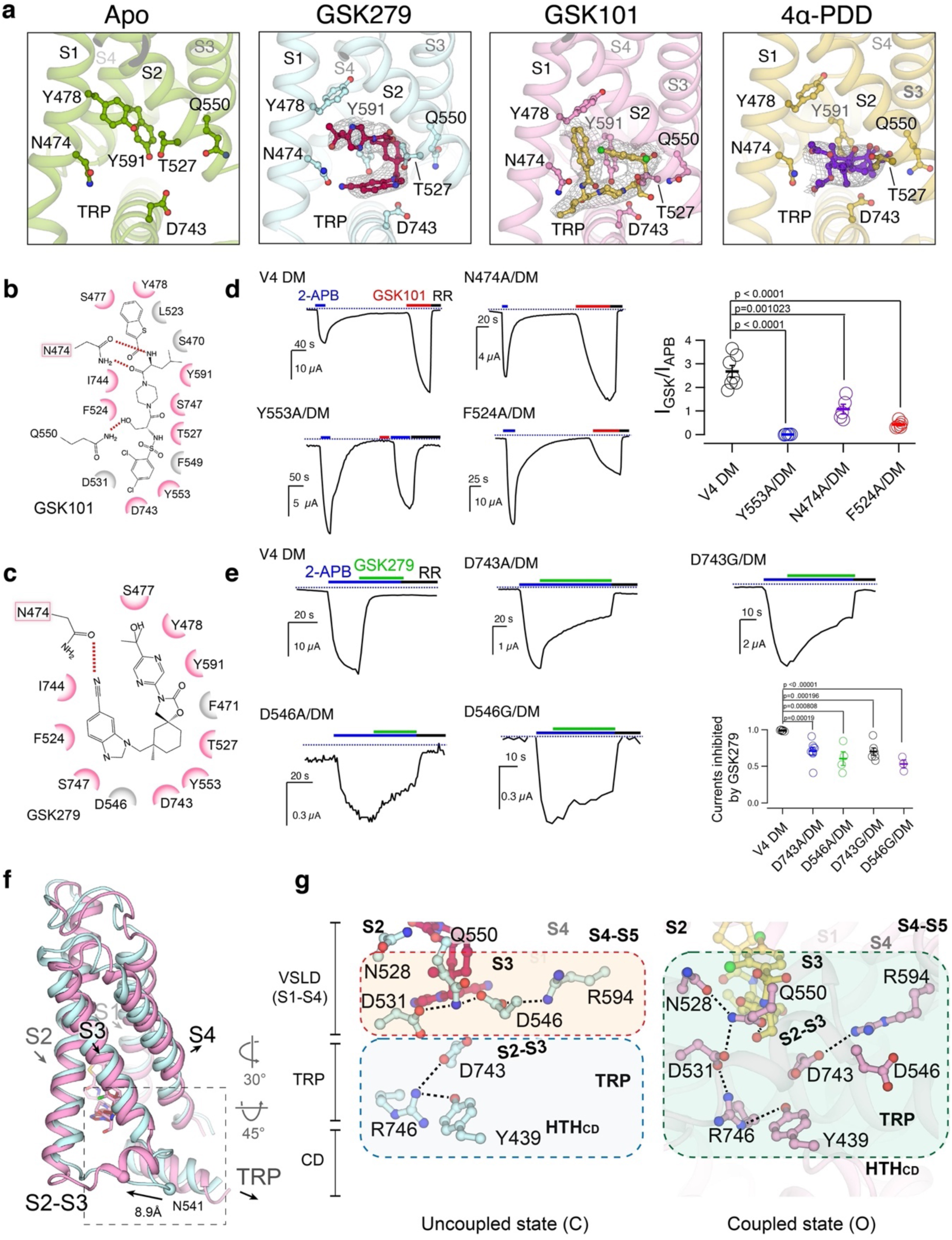
Antagonist- and agonist-binding in the hTRPV4 channel. **a**, Cryo-EM densities (gray mesh) for ligands (GSK279; red stick, GSK101; gold stick, and 4α– PDD; violet stick) in apo, closed, open, and 4α–PDD-open. Densities are contoured at 0.23, 0.36, 0.34, and 0.22 thresholding, respectively. Sidechains of residues are shown in sticks. **b**,**c**, Ligplot schematics of GSK279–TRPV4 and GSK101-TRPV4 interactions, where residues within 3.8 Å to the ligands are shown. Pink colored residues are shared at both interactions. **d**, Probing the agonist binding site. Representative two-electrode voltage clamp (TEVC) recording of human TRPV4 mutant (hTRPV4 DM) and additional mutants made with the background hTRPV4 DM as indicated in the figure. The currents were elicited by 2 mM 2-APB and 5 µM GSK101 as denoted by colored horizontal lines. Voltage was held at -60 mV then ramped to +60 mV over 300 ms every 2 seconds. Plotted here are the currents at -60 mV. The dotted line indicates zero-current level. Right panel: Summary of current responses to 5 µM GSK101 relative to saturating 2-APB (2 mM) at room temperature. Values for individual oocytes are shown as open circles with mean ± S.E.M. shown (n = 5–7). P values are calculated by two-tailed Student’s t test. **e**, Probing the inhibitor binding site. TEVC recordings of hTRPV4 DM (N456H/W737R) and additional mutants made in the background of hTRPV4 DM, as indicated. Currents at -60 mV induced by 2 mM 2-APB then co-application of 4 µM GSK279 followed by 20 µM ruthenium red, as indicated by colored horizontal lines, using the same voltage protocol as in (**c**). Summary of current inhibition by 4 µM GSK279 relative to current from saturating 2-APB (2 mM) at room temperature. Values for individual oocytes are shown as open circles with mean ± S.E.M. shown (n = 3–9). P values are calculated by two-tailed Student’s t test. **f**, Comparison of conformational changes in the VSLD of GSK279-hTRPV4-RhoA (cyan) and GSK101-hTRPV4-RhoA (pink). Ligands are shown as sticks. Arrows indicate helix movements and rotations. **g**, Comparison of coupling networks at the VSLD, CD and TRP domain in closed and open states. Dashed lines indicate hydrogen bonds and salt-bridges.

How is it that GSK279 and GSK101 impose opposite effects on TRPV4 gating while binding to a common set of residues in the VSLD cavity? We observe that going from the GSK279-bound closed state to the GSK101-bound open state, the S2-S3 linker (M534-S548) undergoes a loop-to-helix transition, thereby altering the interaction network among the VSLD, TRP domain, and CD (Fig. 2f). In the closed state, D531 (S2), Q550 (S3), D546 (S2-S3), and R594 (C-terminal half of S4; S4b) form a charged H-bond relay within the VSLD, while D743, R746 (TRP domain) and Y439 (CD; HTH_CD_) connect the TRP domain and the CD (Fig. 2g, left). These two separate interaction networks appear to be decoupled between the VSLD and the TRP/CD. In contrast, the rearrangement of the S2-S3 linker in the open state couples the VSLD to the TRP domain and the CD. First, the D531-Q550-D546-R594 interactions within the VSLD are broken. As a result, Q550 (S3) interacts with both the hydroxyl group of GSK101 and N528 (S2), while R594 (S4b) forms a new salt-bridge interaction with D743 (TRP domain), and D531 (S2) interacts with R746 (TRP domain) (Fig. 2g, right), resulting in the ∼4 Å downward swing of the TRP domain and rearrangement of the CD (HTH_CD_) (Figs 2f,g and Extended Data Figs. 5b,c). Therefore, the opposing actions of agonist/antagonist binding to the same cavity in TRPV4 originate from the coupling or decoupling of the VSLD-TRP-CD subdomains mediated by ligand binding (Fig. 2g).

These conformational changes associated with channel activation propagate from the ligand-binding site to the pore domain. The S4-S5 linker, S5, and PH rotate toward the central ion conduction pathway, and S6 undergoes changes in the π-helix position (from M718 to V708) and rotation at the C-terminus (C-terminal half of S6 [S6b]; residues N712-G719) (Extended Data Fig. 5b).

### Pore conformation changes during TRPV4 gating

We observe extensive conformational changes in the TRPV4 pore domain during ligand-dependent gating. The GSK279-bound closed state adopts a wide-set SF (I678-G679-M680) and the narrowest constriction point of the S6 gate is at M718 (Figs. 3a-c). The diagonal distance between G679 backbone carbonyls in the SF (12.2 Å) is too far to coordinate cations, while that between the M718 sidechains (4.9 Å) is too narrow for ion conduction. However, in the GSK101-bound open state, the SF and the PHs move closer to enable cation coordination (the diagonal distance of G679 backbone carbonyl is 7.0 Å), which is consistent with previous mutagenesis studies^33^. Meanwhile the pore-lining S6 helices rotate ∼90°, thus changing the gate position (from M718 to I715) and the gate opening (the diagonal distance at I715 is 8.4 Å) (Figs. 3c,d). This S6 rotation between the two states is due to the shift of the π-helix position on S6 (Fig. 3a), resulting in a local secondary structure conversion between α- and π-helix along S6b, which is unique amongst the TRP channels^34^. The conformational changes of S6 are facilitated by rearrangement in the interfacial contacts with the neighboring S4-S5 linker (Extended Data Fig. 5d).

**Fig. 3.**
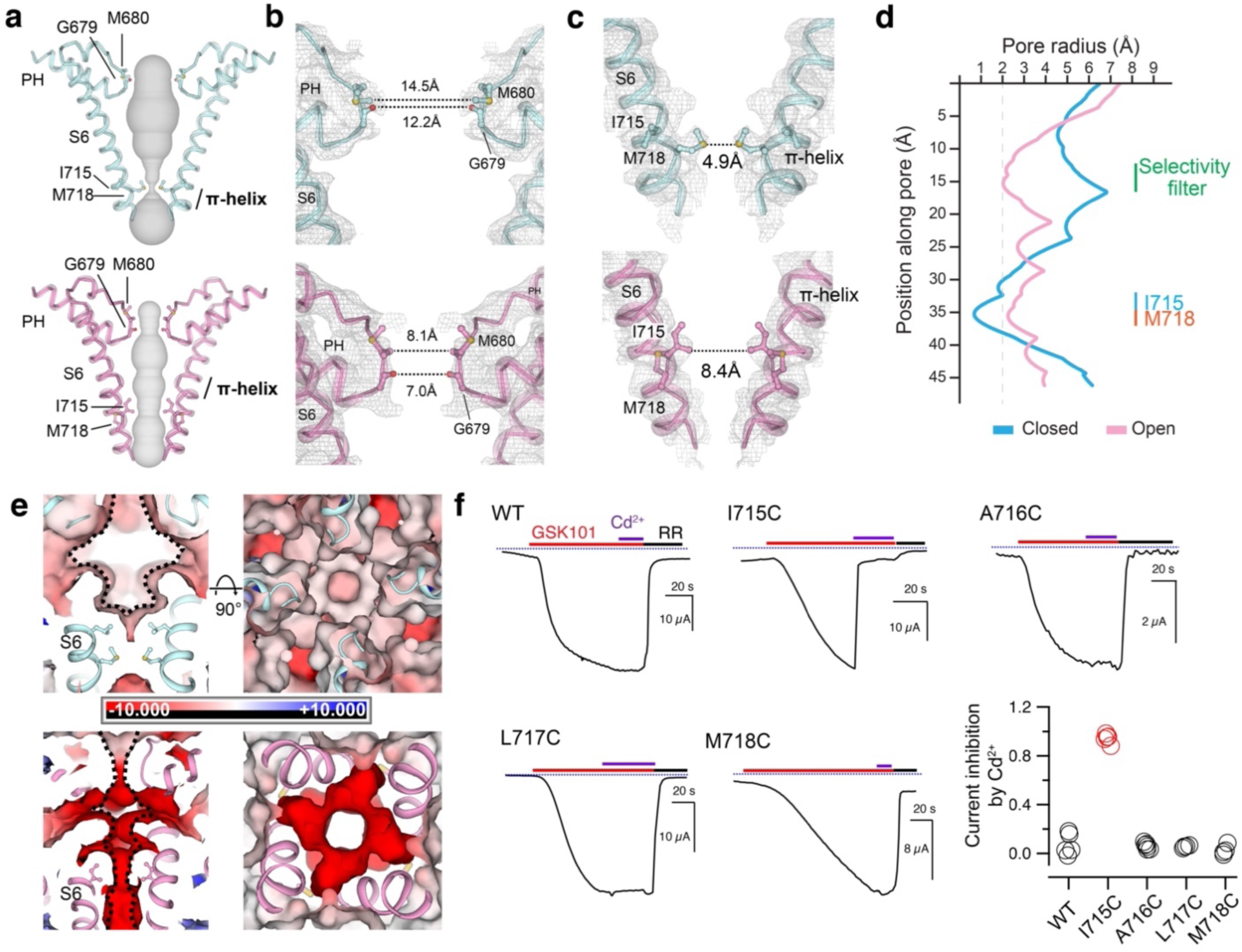
Structural changes in the S6 gate and SF position during channel gating. **a**, Ion permeation pathway in the closed and open state shown as gray surfaces. S6 helices from two protomers are shown in cartoon. Only two subunits are shown for clarity. Gating and selectivity filter residues shown as sticks. **b**,**c**, Close-up views of the SF region (**b**) and S6 gate (**c**) for the closed and open states. The dotted lines indicate diagonal distances between gating residues of opposite protomers. Gray mesh indicates cryo-EM densities contoured at 0.3 (top) and 0.4 (bottom) thresholding, respectively. **d**, Pore radii calculated using the HOLE program in Coot for representative TRPV4 structures as color-coded. The minimal radius for a hydrophobic gate to be open is considered 2.0 Å. Residues corresponding to the SF (M680 and G679) and the S6 gate (I715 and M718) are denoted. **e**, APBS surface electrostatics of the pore in the closed and open states as viewed from the membrane plane (right) and from the extracellular side (left). S6 helices and SF region are shown in cartoon and gating residues as sticks. **f**, Representative time-course recording of hTRPV4 WT and mutants. Currents elicited by 5 µM GSK101 and co-application with 10 µM Cd^2+^ followed by 20 µM ruthenium red (RR) as indicated by colored horizontal lines. The voltage was ramped from -60 mV to +60 mV in 300 ms every 2 seconds. The currents at -60 mV were used for the plot. Dotted blue lines indicate zero-current level. Right panel, summary of current inhibition by 10 µM Cd^2+^ relative to 5 µM GSK101 induced currents. Values for individual oocytes are shown as open circles (n = 4–7).

Interestingly, when compared to the closed state, we found that the pore cavity volume is reduced, and the surface electrostatic potential becomes substantially more negative in the open state (Fig. 3e). This was not observed in other TRPV channels^28^. Taken together, we posit that these unusual changes in the pore help form the ion permeation path, similar to the canonical tetrameric cation channel^28^.

To validate the S6 gate position observed in the open-state structure, we performed Cd^2+^-dependent blocking of TRPV4. Extracellular application of Cd^2+^ does not affect the opening of WT TRPV4 by GSK101 (Fig. 3f). Among cysteine mutants near the S6 gate (I715C, A716C, L717C, and M718C), only I715C showed substantial inhibition of GSK101-elicited current by extracellular Cd^2+^, supporting the observation that I715 faces the ion permeation pathway in the TRPV4 open state (Fig. 3f).

### Neuropathy mutations disrupt RhoA binding to TRPV4

At the core of the small GTPase RhoA is a highly conserved G domain comprising a six-stranded β-sheet (β1-β6), six helices (α1-α6), and a C-terminal variable region (Fig. 4a and Extended Data Fig. 6). Switch I and II regions within the G domain adopt distinct GDP- or GTP-dependent conformations, where the latter favors effector protein binding^35^.

**Fig. 4.**
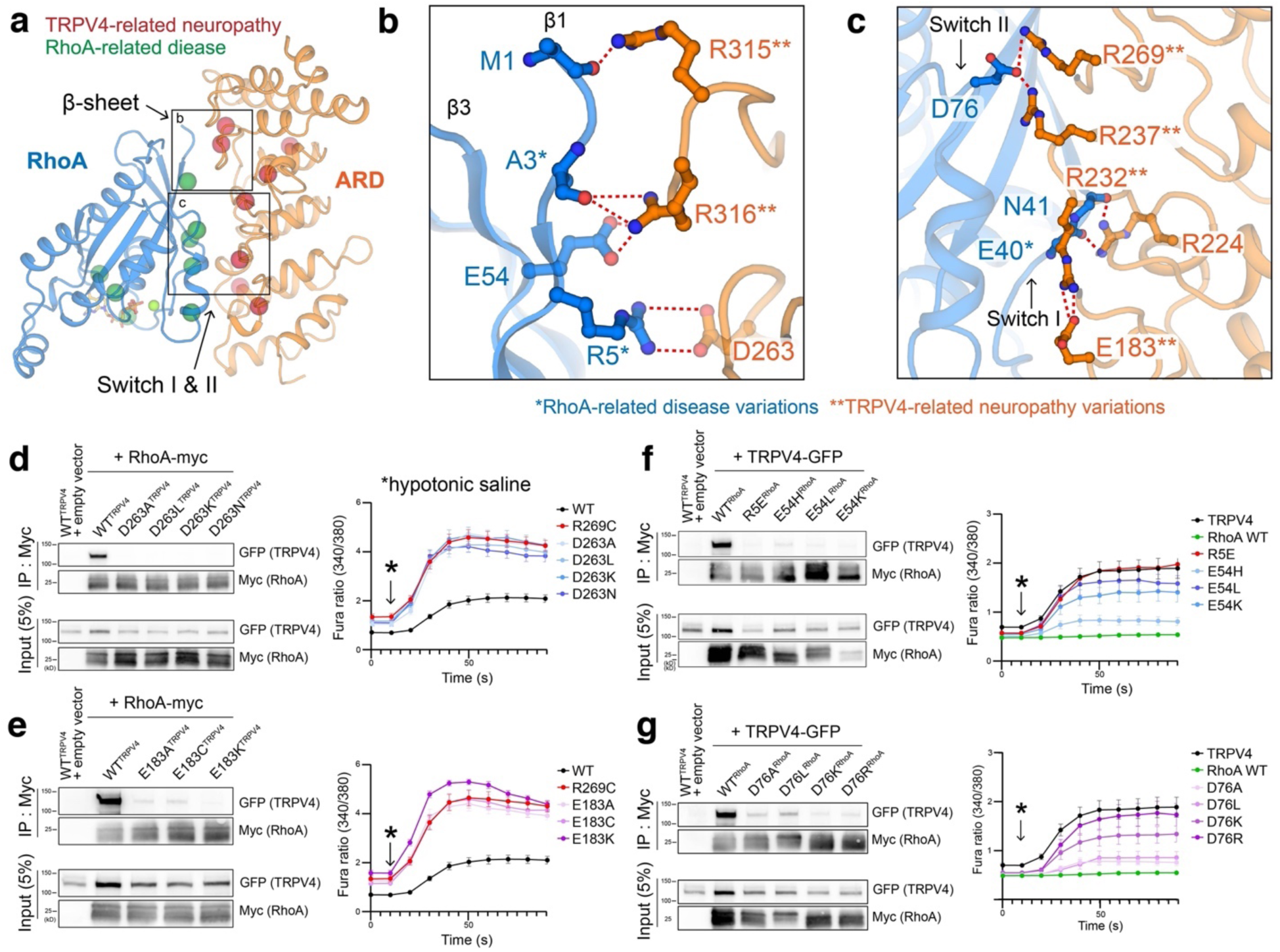
Interaction between hTRPV4 ARD and RhoA. **a**, Overall interaction interface of hTRPV4 ARD and RhoA. Disease-causing mutations mapped onto the TRPV4-RhoA interface. Mutations causing peripheral neuropathy of TRPV4- and RhoA-related diseases are shown as red and green spheres, respectively. **b**,**c**, Detailed hTRPV4 ARD – RhoA interactions within the β1-β3 region (**b**) and switch region (**c**). RhoA residues are colored blue, TRPV4 residues are colored orange. * disease-causing mutations in RhoA and ** neuropathy-causing mutations in TRPV4. The red dashed lines indicate salt-bridge interactions. **d-g**, (Left panels) Co-immunoprecipitation of HEK293T cells transfected with TRPV4-GFP (**d**, E183A/C/K; **e**, D263A/L/K/N) and RhoA-Myc (**f**, R5E, E54L/H/K; **g**, D76A/L/K/R) demonstrates that mutations at the TRPV4-RhoA interface reduces their interaction. (Right panels) Representative plots from ratiometric calcium imaging experiments. MN-1 cells were transfected with GFP-tagged TRPV4 plasmids only (**d** and **e**) and GFP-tagged TRPV4 and RhoA plasmids (**f** and **g**) and loaded with Fura-2 AM calcium indicator. Baseline and hypotonic-stimulated calcium responses were then measured over time.

We performed focused-refinement on the RhoA and ARD region in the 3D reconstructions of the GSK279-bound, apo, and GSK101-bound data. Although the EM density for the ARDs in all three states exhibited high quality, that for RhoA was most well resolved in the GSK279-bound closed state, which we used to unambiguously model RhoA and map its interaction with TRPV4 (Extended Data Figs. 2, 3a, and 3b). We resolved GDP within the RhoA-TRPV4 signaling complex in the GSK279-bound closed state, and by including GTPγS before freezing grids we observed density consistent with GTPγS in the open state (Extended Data Figs. 7a,b). We did not, however, observe substantial structural differences of RhoA between the GDP- and GTPγS-bound forms in complex with the closed- and open-states of TRPV4, respectively (Extended Data Fig. 7c). We used the model from the GSK279-bound closed state for our analysis of RhoA-TRPV4 interactions. Notably, while the switch I in RhoA resembles the GDP-bound RhoA structure, the switch II is distinct from either the GDP- or the GTP-bound RhoA structure (PDB IFTN and 1A2B; Extended Data Fig. 7d). The interfacial contact that associates TRPV4 and RhoA is mediated principally by electrostatic interactions between β1 and β3, switch I, and switch II in RhoA and AR2-AR5 in TRPV4 (Fig. 4a and Extended Data Fig. 7e). Sequence comparisons indicate that the interfacial residues are unique to the TRPV4 ARD across all TRPV channels, and to Rho isoforms amongst other small GTPases, suggesting that the observed RhoA binding mode is specific to TRPV4 (Fig. 6)^22^.

Strikingly, most residues mutated in TRPV4-mediated neuropathy (R237, R269, R315, and R316) are clustered at the interface with RhoA (Fig. 4a). R232, a reported TRPV4-neuropathy mutation site, does not directly participate in the interfacial contact but does form an intra-subunit salt-bridge with another neuropathy-causing residue E183, which is likely important for tuning electrostatics and the local conformation of the RhoA-binding surface of the ARD (Extended Data Fig. 7f). Therefore, disease-causing mutations in TRPV4 likely weaken the interactions with RhoA^22^. Notably, we also found that several cancer-related mutation sites in RhoA, such as A3, R5, and E50^36–38^, are located at the interface with the TRPV4 ARD (Figs. 4b,c), providing evidence for an interplay between TRPV4 and RhoA in cancer^5,21,39^, although these RhoA mutations may also impact effector binding more broadly.

To probe the interactions between TRPV4 and RhoA, we first performed co-immunoprecipitation of GFP-tagged WT or mutant TRPV4 with Myc-tagged WT or mutant RhoA overexpressed in HEK293T cells^22^. We previously showed that neuropathy-causing mutations (R232C, R237L, R269C, R315W) in TRPV4 disrupt the interaction with RhoA, consistent with our structures (Figs. 4b,c). We further mutated residues in RhoA (R5, E54, D76) and TRPV4 (E183 and D263) that form the RhoA-TRPV4 interface. All mutants tested (R5E^RhoA^, E54H/L/K^RhoA^, bbbbbbh ncD76A/L/K/R^RhoA^, E183A/C/K^TRPV4^, and D263A/L/K/N^TRPV4^) substantially decreased the amount of immunoprecipitated partner proteins (TRPV4 and RhoA) (Fig. 4d). We then tested the effects of these mutations on TRPV4 function using ratiometric calcium imaging. When we tested TRPV4 mutant function, we opted not to overexpress RhoA but rather to rely on interaction with endogenous RhoA. Compared to WT TRPV4, E183A/C/K or D263A/L/K/N mutations result in increased basal and hypotonic saline-induced calcium influxes, similar to the neuropathy mutant R269C mutation^22^. We then tested the effect of RhoA mutations on TRPV4 channel activity, in experiments in which we overexpressed both TRPV4 and RhoA. Whereas expression of WT RhoA strongly suppressed both basal and hypotonic saline induced Ca^2+^ influx (Figs. 1b and 4d), RhoA mutants demonstrated reduced suppression of TRPV4 channel activity (Fig. 4d). Importantly, the degree of suppression of TRPV4 activity correlated with the interaction strength between the RhoA mutants and TRPV4 (Fig. 4d). These data demonstrate that TRPV4-RhoA interaction strength strongly correlates with TRPV4 channel activity, and that disruption of this interaction underlies the gain of function due to neuropathy mutations within the ARD. This suggests that balancing the two different neurophysiological signaling pathways (calcium signaling and actin cytoskeleton remodeling) requires fine tuning of the TRPV4-RhoA interaction.

### RhoA-dependent TRPV4 gating

Despite only subtle conformational changes in RhoA, the buried surface area between RhoA and TRPV4 in the open state is reduced (∼684 Å^2^) compared to that observed in the closed state structure (∼752 Å^2^), suggesting RhoA-TRPV4 interactions become weaker in the open state (Extended Data Fig. 7g). To compare the relative occupancy and/or dynamics of RhoA bound to TRPV4 in different channel functional states, we lowpass filtered the cryo-EM maps of the closed, apo, and open state structure at the same resolution (4 Å) and applied the same value of B-factor sharpening. Notably, the EM density for RhoA progressively decreases from the closed to the apo then to the open states, suggesting that the occupancy decreases and/or bound RhoA becomes more flexible during TRPV4 gating (Fig. 5b). Consistent with this observation, further cryo-EM 3D classifications of the closed and the open state reconstructions show that the closed state reconstruction contains a major class with strong RhoA density while the open state reconstruction contains classes with weak RhoA density (Extended Data Fig. 9). The apparent differential resolution of RhoA density between the closed, apo, and open states led us to hypothesize the state-dependent RhoA binding affinity to TRPV4.

**Fig. 5.**
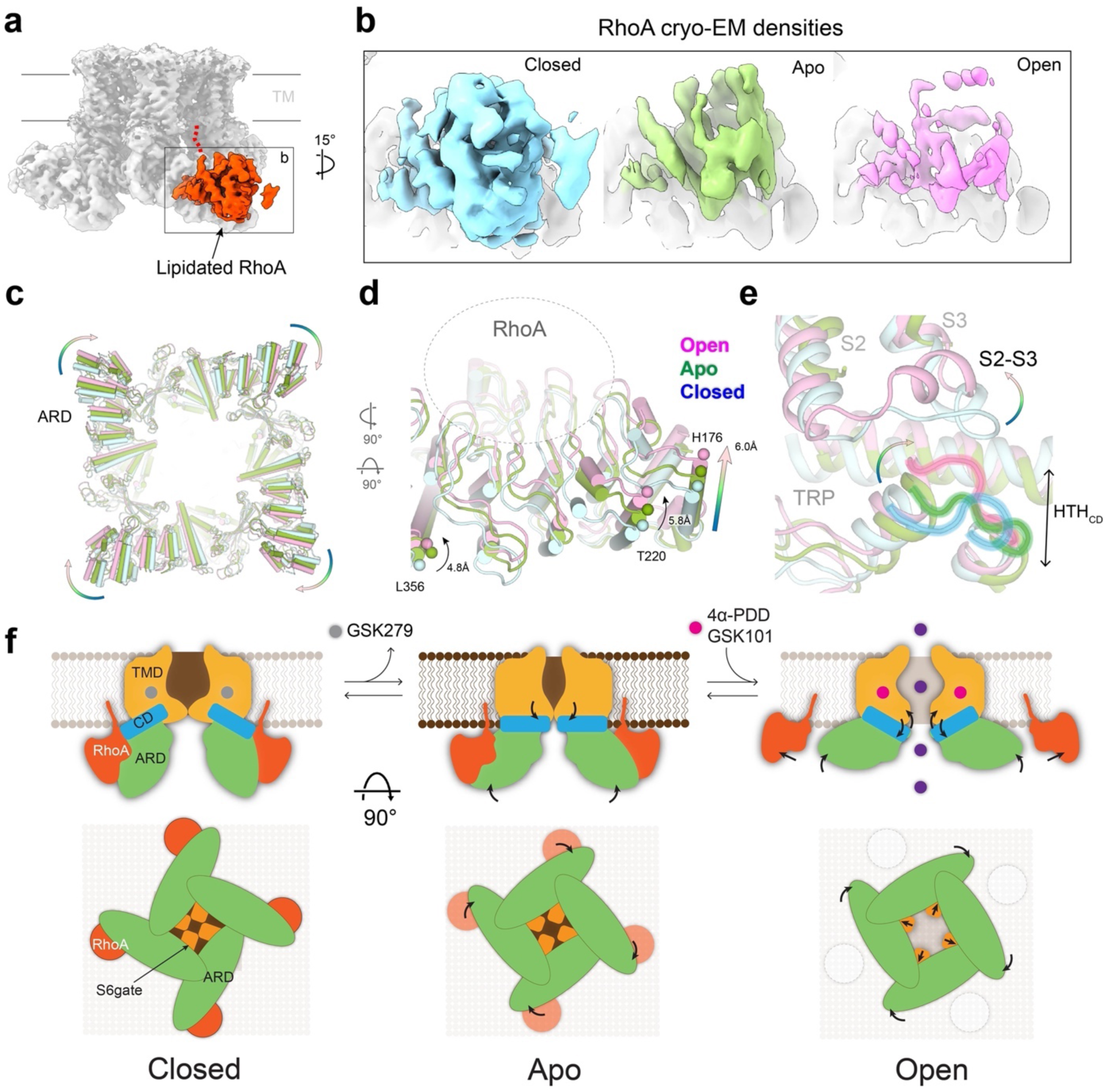
Structural basis of RhoA-dependent gating of hTRPV4. **a**, Cryo-EM reconstruction of GSK279-hTRPV4-RhoA, with the RhoA from one protomer highlighted in red at thresholding 0.29. **b**, Side-by-side comparison of cryo-EM densities of RhoA in closed (cyan), apo (green), and open (pink) states at thresholding 0.25. **c-e**, Comparison of closed, apo, and open structures viewed from the intracellular side (**c**). Close-up view of the ARD (**d**). and coupling domain (**e**; HTH_CD_, TRP, and S2-S3). Arrows indicate movements of the ARD. ARD/CD movement occurs at an individual protomer level. **f**, Ligand-dependent channel gating of hTRPV4–RhoA. Schematic illustration of the conformational rearrangements in the S6 gate, ARD, and RhoA during TRPV4 gating.

TRPV channels are highly allosterically coupled across their domains, as was previously shown^40^, and because we observed progressive changes in RhoA occupancy during gating we attempted to infer the effect of RhoA on TRPV4 gating. Structural alignment at the TMDs show that each ankyrin repeat (AR) in the ARD swings toward the membrane by ∼6 Å from the closed to the open state (Fig. 5d). When viewed from the cytoplasmic side, individual ARs rotate clockwise in addition to the upward swing of the ARD, rather than a concerted rotation of the tetrameric ARD ring (Fig. 5c). Movements in the ARD propagate as conformational changes in the CD (HTH_CD_), and in turn the TRP domain and the VSLD (Fig. 5e), as essential gating steps in TRPV4 activation (Fig. 2). RhoA, which is anchored to the membrane bilayer via its prenylated C-terminus and binds to TRPV4 at the cytoplasmic ARD, exerts forces on the ARD analogous to a clamp, which suppresses ARD motion and thus TRPV4 activation (Fig. 5f).

## Discussion

Here we resolve, for the first time, human TRPV4 structures in complex with the small GTPase RhoA and elucidate structural mechanisms underlying the crosstalk between RhoA and TRPV4. The rigid-body rotation of the ARD domain couples the CD, the TRP domain, and VSLD, leading to TRPV4 opening (Figs. 2g and 5e). Through membrane-anchored RhoA binding/dissociation, Nature has designed a means to control the ARD rearrangement and therefore TRPV4 gating. Although our current studies focus on the effect of RhoA on TRPV4 function, conversely, TRPV4 activity can modulate RhoA unbinding and release^22^. Once TRPV4 is activated, the membrane-bound RhoA is released to regulate cytoskeleton dynamics upon activation by TRPV4-mediated calcium influx^41^.

While the localization of neuropathy-associated TRPV4 mutations at the TRPV4-RhoA interface suggests that control of this signaling complex is particularly important in the nervous system, TRPV4-RhoA interactions are likely to play a fundamental role in signaling in other disease states. Intriguingly, we demonstrated that several cancer-related mutations of RhoA also disrupt the TRPV4-RhoA binding interface. Although these mutations may interrupt RhoA binding with several partners, these data are consistent with prior work demonstrating a role for TRPV4-RhoA in the metastatic cascade and tumor vascularization^5,21,39^. TRPV4-RhoA interactions are also likely essential in endothelial (e.g. lung, retinal, intestinal) and epithelial (e.g. skin, lung) cells, where both proteins have been shown to play key roles in barrier maintenance ^21,39,42^. Importantly, numerous studies have reported that RhoA and other small GTPases can affect the function of various ion channels, including K^+^ channels, several other TRP channels, epithelial Na^+^ channels, and acid-sensing ion channels^12–16^. Further work will be needed to determine whether these other functional channel-GTPase couplings involve direct interactions, and if so, whether similar or distinct structural mechanisms are utilized to those described here.

TRPV4 is a well-established osmosensor^18^ and is also reported to respond to shear stress, but the structural mechanisms mediating channel activation by these stimuli have not been determined. Osmosensors can sense either the direct change in the extracellular water activity or subsequent modifications in cellular organization^43^. We speculate that the wide-set pore domain in the closed state of TRPV4 and its unusual pore conformational changes upon channel activation contribute to channel sensitivity to changes in extracellular fluid activity or shear stress. Moreover, the direct association with plasma membrane-anchored RhoA may enable TRPV4 to detect changes in cellular shape and morphology induced by osmotic shock or mechanical force. Consistent with this speculation, RhoA is activated under cell stretch^44^, and the interaction of TRPV4 with actin is essential for cell-swelling-induced channel activation^45^.

Our studies also deciphered mechanisms of ligand-dependent TRPV4 activation and inhibition. The binding of antagonists and agonists to a shared site within the VSLD cavity leads to opposing conformational changes on the TRPV4 pore. Our structural analysis elucidated that agonist binding couples, whereas antagonist binding decouples, the interaction network amongst the VSLD, the TRP domain, and the CD. Our discoveries of the human TRPV4 structure bound to the clinical candidate antagonist GSK279, as well as the molecular mechanisms of ligand-dependent TRPV4 gating will facilitate the development of drugs targeting TRPV4. For example, with this structure, one can modify GSK279 or find compounds with new scaffolds (via virtual screening) to increase their potency or efficacy. Furthermore, the presented structures delineate principles of channel inhibition that can be utilized for virtual screening for compounds that could bind TRPV4 at different locations but exert similar inhibitory decoupling effects.

**Table 1.** Cryo-EM data collection, refinement, and validation statistics.

|  | hTRPV4-RhoA<br>GSK2798745<br>(EMDB-28975)<br>(PDB-8FC7) | hTRPV4<br>GSK1016790A<br>(EMDB-28976)<br>(PDB-8FC8) | hTRPV4-RhoA<br>GSK1016790A<br>(PDB-8FCB) | hTRPV4-RhoA<br>apo<br>(EMDB-28977)<br>(PDB-8FC9) | hTRPV4<br>4α-PDD<br>(EMDB-28978)<br>(PDB-8FCA) |
| --- | --- | --- | --- | --- | --- |
| **Data collection and processing** |  |  |  |  |  |
| Magnification | 81,000 | 81,000 |  | 81,000 | 81,000 |
| Voltage (kV) | 300 | 300 |  | 300 | 300 |
| Electron exposure (e<sup>-</sup>/Å<sup>2</sup>) | 60 | 60 |  | 60 | 60 |
| Defocus range (μm) | -0.8 to -1.8 | -0.8 to -1.8 |  | -0.8 to -1.8 | -0.8 to -1.8 |
| Pixel size (Å) | 1.08 | 1.08 |  | 1.08 | 1.08 |
| Symmetry imposed | C4 | C4 |  | C4 | C4 |
| Initial particles images (no.) | 2,111,563 | 8,119,132 |  | 3,412,804 | 4,418,471 |
| Final particles images (no.) | 119,462 | 207,645 |  | 172,225 | 375,644 |
| Map resolution (Å) | 3.3 | 3.47 |  | 3.75 | 3.41 |
| FSC threshold | 0.143 | 0.143 |  | 0.143 | 0.143 |
| **Refinement** |  |  |  |  |  |
| Initial model used (PDB code) | TRPV1 (7LP9)<br>RhoA (1FTN) | PDB-8FC7 |  | PDB-8FC7 | PDB-8FC7 |
| Map sharpening *B* factor (Å<sup>2</sup>) | -136 | -183 |  | -195 | -140 |
| Model composition |  |  |  |  |  |
| Non-hydrogen atoms | 25,716 | 19,764 | 25,968 | 21,812 | 18,224 |
| Protein residues | 3,284 | 2,468 | 3,228 | 3,092 | 2,352 |
| Ligands | 12 | 8 | 12 | 0 | 4 |
| *B* factors(Å<sup>2</sup>) |  |  |  |  |  |
| Protein | 90.02 | 81.88 | 101.36 | 136.85 | 61.33 |
| Ligand | 83.5 | 46.42 | 68.17 | - | 96.78 |
| R.m.s deviations |  |  |  |  |  |
| Bond lengths (Å) | 0.004 | 0.004 | 0.004 | 0.004 | 0.005 |
| Bond angles (°) | 0.919 | 0.945 | 1.134 | 0.562 | 0.968 |
| **Validation** |  |  |  |  |  |
| MolProbity score | 1.56 | 1.22 | 1.35 | 1.06 | 1.22 |
| Clashscore | 5.17 | 4.06 | 4.48 | 3.81 | 4.12 |
| Poor rotamers (%) | 0 | 0.19 | 0.16 | 0.22 | 0 |
| Ramachandran plot |  |  |  |  |  |
| Favored (%) | 95.83 | 97.88 | 97.38 | 98.56 | 97.59 |
| Allowed (%) | 4.17 | 2.12 | 2.62 | 1.44 | 2.41 |
| Disallowed (%) | 0 | 0 | 0 | 0 | 0 |

## Materials and Methods

### Protein expression and purification

The *Homo sapiens* full-length TRPV4 (hTRPV4) was cloned into a modified pEG BacMam vector^46^ in frame with a FLAG-tag and 10×His-tag at the C terminus. The hTRPV4-RhoA complex was expressed by baculovirus-mediated transduction of the human embryonic kidney (HEK) GnTI^−^ suspension cells, cultured in FreeStyle 293 medium (Life Technologies) with 2% (vol/vol) FBS at 8% CO_2_. hTRPV4 baculovirus was generated following the Bac-to-Bac^®^ Baculovirus Expression System protocol (Life Technologies). Cultures were infected at a cell density of ∼1.5M mL^-1^ with 1.5 – 2 % (v/v) P3 or P4 baculovirus for hTRPV4. After 18 h of shaking incubation at 37 °C, 10 mM sodium butyrate was added, and the growth temperature was lowered to 30 °C to boost protein expression. After 68-72 h, the cells were harvested by centrifugation at 550 xg and resuspended in lysis buffer (20 mM Tris pH 8, 150 mM NaCl, 12 μg ml^−1^ leupeptin, 12 μg ml^−1^ pepstatin, 12 μg ml^−1^ aprotinin, 2 μg ml^−1^ Dnase I, 1 mM PMSF, 10% (v/v) glycerol, 1 mM dithiothreitol (DTT), 1% (w/v) Lauryl Maltose Neopentyl Glycol (LMNG; Anatrace), and 0.1% (w/v) cholesteryl hemisuccinate (CHS; Anatrace). Membranes were solubilized at 4 °C by gentle agitation for 2 h followed by centrifugation at 8,000xg for 30 min to remove insoluble material. The supernatant was subsequently incubated with anti-FLAG M2 resin (Sigma-Aldrich) for 40 min at 4 °C with gentle agitation. The resin was then packed onto a gravity-flow column (Bio-Rad) and washed with 10 column volumes (CV) of high salt wash buffer (20 mM Tris pH 8.0, 150 mM NaCl, 0.1% LMNG, 0.01% CHS, 5mM ATP, 1 mM DTT, and 5% glycerol) followed by 10 CV of low salt wash buffer 1 (20 mM Tris pH 8.0, 500 mM NaCl, 0.1% LMNG, 0.01% CHS, 1 mM DTT, and 5% glycerol), and then 10 CV of low salt wash buffer 2 (20 mM Tris pH 8.0, 150 mM NaCl, 0.03% LMNG 0.003% CHS, 1mM DTT and 5% glycerol). The hTRPV4-RhoA complex was eluted by 5CV of elution buffer (20 mM Tris pH 8.0, 150 mM NaCl, 0.03% LMNG, 0.003% CHS, 1 mM DTT, 5% glycerol, 0.150 mg mL^-1^ FLAG peptide). The eluted protein complex was concentrated and further purified by size exclusion chromatography (SEC) on a Superose 6 Increase column (Cytiva Life Science) equilibrated with SEC buffer (20 mM Tris pH 8.0, 150 mM NaCl, 0.00075% LMNG, 0.000075% CHS, 0.0003% glycol-diosgenin (GDN; Anatrace), 1 mM DTT, and 5% glycerol).

### Cryo-EM sample preparation and data acquisition

hTRPV4-RhoA peak fractions from SEC were concentrated to 0.8 – 1.2 mg mL^-1^. All samples were incubated with different ligand conditions at 4 °C for 15-20 min prior to freezing grids. For the apo-hTRPV4-RhoA sample, 2% (v/v) DMSO was added to the purified protein instead of ligands. For the GSK279-hTRPV4-RhoA, sample was incubated with 20 μM GSK2798745 (GSK279; Medchemexpress). For the GSK101-hTRPV4-RhoA, 20 μM GSK1016790A (GSK101; Tocris) and 2 mM Guanosine 5’-[γ-thio] triphosphate (GTPγS; Sigma-Aldrich) were incubated with the protein. For the 4α-PDD-hTRPV4-RhoA, sample was incubated with 4α-Phorbol 12,13-didecanoate (4α-PDD; Sigma-Aldrich) and 2mM GTPγS. All grids were prepared with a Leica EM GP2 plunge freezer (Leica) at 4 °C and 95% humidity. 3 μL of sample was applied to a freshly glow-discharged UltrAuFoil R1.2/1.3 300 mesh grids (Quantifoil), blotted for 1.5 – 2.0 s, depending on the specific ligand conditions, to obtain optimal ice thickness for data collection.

Cryo-EM datasets for the apo-hTRPV4-RhoA, GSK279-hTRPV4-RhoA, GSK101-hTRPV4-RhoA, and 4α-PDD-hTRPV4-RhoA were collected with Titan Krios microscope (FEI) operating at 300 keV equipped with a K3 detector (Gatan) with GIF BioQuantum energy filter (20 eV slit width: Gatan) in counting mode, using the Latitude-S automated data acquisition program (Gatan). Movie datasets were collected at a nominal magnification of 81,000x with a pixel size of 1.08 Å per pixel^-1^ at specimen level. Each movie contained 60 frames over 4.6 s exposure time, using a dose rate ∼15 e^-^Å^-2^s^-1^, resulting in the total dose of ∼60 e^-^Å^-2^. The nominal defocus ranged from -0.8 to −2.0 μm.

### Cryo-EM data processing

A total of 5,666, 3,877, 17,040, and 8,226 movies were collected for the apo-hTRPV4-RhoA, GSK279-hTRPV4-RhoA, GSK101-hTRPV4-RhoA, and 4α-PDD-hTRPV4-RhoA structures, respectively. All four datasets were processed similarly, as illustrated in Extended Data Fig. 1c with RELION 4.0^47^ and cryoSPARC^48^. Beam-induced motion correction and dose-weighting were performed using MotionCor2^49^, followed by CTF estimation using Gctf^50^ in RELION. Micrographs were subsequentially selected based on astigmatism, CTF fit quality, CTF estimated maximum resolution, and defocus values. An initial set of particles were manually picked and subjected to a reference-free 2D classification (k = 7, T = 2), from which the best two to three classes were selected as reference for a templated-based auto-picking in RELION. Particles were re-centered, and re-extracted Fourier binned 4×4 (64-pixel box size) and imported to cryoSPARC. Two rounds of 2D classification with 50 classes were used to remove contamination and false noise picks, like chaperones and junk particles. The particles were imported to RELION and subjected to reference-free 2D classification (k = 50, T = 2), ignoring the CTFs until the first peak option. Classes were selected that showed apparent structure features of the hTRPV4-RhoA complex. These particles were subsequently subjected to 3D classification (k=3 or 4, T=8) with C1 symmetry with image alignment using a previously published TRPV1 map (EMD-23473, low-passed filtered to 60 Å) as a reference without masking. The class showing the apparent shape of the hTRPV4-RhoA complex was selected, re-centered, and re-extracted Fourier binned 2×2 for further classification. Refined particles at 2×2 Fourier binning were subjected to 3D classification with image alignment (K=3, T=8) and C1 symmetry. The class showing substantial densities for transmembrane helices and RhoA, was selected, re-centered, re-extracted without binning, and subjected to 3D classification without image alignment (k=2 or 3, T= 8-12) with a soft mask covering TRPV4 and RhoA with C4 symmetry imposed. The particles from the class with the best-resolved transmembrane helices were subjected to 3D-auto-refinement with a soft TRPV4-RhoA mask. The refined particles were processed for CTF refinement,^51^ Bayesian polishing^52^ then subjected to particle subtraction to resolve the strong density at the transmembrane domains (TMDs), followed by focused 3D classification. The tight mask for signal subtraction was made by subtracting out all unnecessary signals from the previous consensus 3D reconstruction, including detergent micelle, parts of the cytosolic domains, and RhoA. The subtracted particles by the tight mask were subjected to focused-3D classification without image alignment (k = 2, T =16 or 20). Particles comprising the best-featured class at TMDs were reverted to original particles, which were input to 3D auto-refinement with a hTRPV4-RhoA full mask. Additional CTF refinement and Bayesian polishing were performed to improve the overall map quality. Finally, particles yielding the best 3D reconstruction from RELION were imported into cyroSPARC and subjected to non-uniform (NU) refinement^53^ with a full mask. To improve the EM density quality at the TMDs of TRPV4 the particles were subjected to particle subtraction of the four RhoA densities, followed by NU refinement. In order to resolve the unambiguous EM density around RhoA for GSK279-hTRPV4-RhoA and GSK101-hTRPV4-RhoA structures, we performed local refinement at the level of one monomeric ARD and RhoA by subtracting signals of the tetrameric TMDs, the other three ARDs, and RhoAs. A detailed data processing flowchart for the GSK101-hTRPV4-RhoA structure is illustrated in Extended Data Fig. 2.

### Model building, refinement, and validation

For manual model building in Coot^54^, the published cryo-EM structure of *Rattus norvegicus* TRPV1 (PDB: 7LP9) and published crystal structure of GDP-bound RhoA (PDB: 1FTN) were docked into the cryo-EM map for GSK279-hTRPV4-RhoA. The angle between the ARD and TMD was first rigid body adjusted into the EM densities, separately. Secondary structures were then rigid body fit into the EM densities using bulky aromatic residues to ensure correct register assignment.

The focus refined TRPV4 channel map and ARD-RhoA focused map facilitated register assignment at S4-S5, S6-TRP flexible linkers and RhoA (Extended Data Fig. 3). The placement of individual residues was adjusted by rigid body fitting then manually refined using real space refinement in Coot, with ideal geometric and secondary structure restraints. The GSK279-hTRPV4-RhoA models served as the initial reference for model building of the GSK101-hTRPV4-RhoA, apo-hTRPV4-RhoA, and 4α-PDD-hTRPV4-RhoA structures. The restraints for ligands and lipid, including GSK279, GSK101, 4α-PDD, and CHS were generated from isomeric SMILES strings using the eLBOW tool^55^ in PHENIX to fix bond lengths and angles. The manually built structure models with ligands were subjected to real-space refinement in PHENIX using cryo-EM maps with global minimization, rigid body refinement and B-factor refinement^56^.

Problematic regions identified by the MolProbity server(http://molprobity.biochem.duke.edu)57, including geometry outliers and Ramachandran outliers, were manually adjusted in Coot. The FSC curves for the model against the full map and both half-maps were generated by comprehensive validation^58^ in PHENIX. The FSCs were in good agreement with each other, indicating models were not over-fitted and refined. Structural illustrations and analysis were performed in Coot^54^, PyMO^59^, UCSF Chimera^60^, and UCSF ChimeraX^61^. For the figure preparations of cryo-EM density maps, UCSF ChimeraX and PyMOL were used.

### Two-electrode voltage-clamp electrophysiology

The WT human TRPV4 DNA was subcloned into the pGEM-HE vector, the construct was linearized with NheI, and complementary RNA (cRNA) was synthesized by in vitro transcription using T7 RNA polymerase (Thermo Fisher). All defolliculated oocytes were purchased from Ecocyte (Austin, TX). Oocytes were injected with cRNA for each of the constructs and incubated at 17°C for 1-3 days in a solution containing 96 mM NaCl, 2 mM KCl, 1 mM MgCl2, 1.8 mM CaCl2, 5 mM HEPES, pH 7.6 (with NaOH), and gentamicin. For the two-electrode voltage-clamp recording, oocyte membrane voltage was controlled using an OC-725C oocyte clamp (Warner Instruments). Data were filtered at 1–3 kHz and digitized at 20 kHz using pClamp software (Molecular Devices) and a Digidata 1440A digitizer (Axon Instruments). Microelectrode resistances were 0.1–1 MΩ when filled with 3 M KCl. The external recording solution contained 100 mM KCl, 2 mM MgCl_2_, 10 mM HEPES, pH 7.6 (with KOH). Agonists, antagonist, and ruthenium red were applied using a gravity-fed perfusion system that can exchange the 150 µL recording chamber volume within a few seconds.

### Antibodies and reagents

Primary antibodies used were rabbit anti-Myc (Cell Signaling Technology, 2272), mouse anti-Myc (Cell Signaling Technology, 2276), rabbit anti-GFP (Thermo Fisher Scientific, A-11122), mouse anti-GFP (Thermo Fisher Scientific, A-11120), rabbit anti-RhoA (Cell Signaling Technology, 67B9), Secondary antibodies used were HRP-conjugated monoclonal mouse anti-rabbit IgG, light chain specific (Jackson ImmunoResearch, 211-032-171) and goat anti-rabbit IgG (Li-COR, 926-32211)

### Co-immunoprecipitation

HEK293T cells were cultured in Dulbecco’s Modified Eagle’s Medium (DMEM) supplemented with 10% (vol/vol) fetal calf serum (FCS) and penicillin/streptomycin at 37°C with 6% CO2. Cells were transfected with Lipofectamine LTX with Plus Reagent (Thermo Fisher Scientific) and lysed 24 h after transfection in IP Lysis Buffer (Pierce, 25 mM Tris-HCl pH 7.4, 150 mM NaCl, 10 mM MgCl2, 1% NP-40, 1 mM EDTA, 5% glycerol) supplemented with EDTA-free Halt protease inhibitor cocktail (Thermo Fisher Scientific). Cells were lysed for 15 min followed by centrifugation at 14,000 RPM for 10 min. Supernatants were incubated with primary antibody bound to magnetic Protein G Dynabeads (Thermo Fisher Scientific) for 1 h at 4°C followed by several washes in IP wash buffer (PBS, 0.2% Tween 20). To elute bound proteins, Laemmli sample buffer with β-mercaptoethanol was added to the beads and samples were heated for 10 min at 70°C. Protein lysates were resolved on 4–15% TGX gels (Bio-Rad Laboratories) and transferred to PVDF membranes. Membranes were developed using SuperSignal West Femto Maximum Sensitivity Substrate (Thermo Fisher Scientific) and imaged using an ImageQuant LAS 4000 system (GE Healthcare).

### Calcium imaging

MN-1 cells were transfected with TRPV4-GFP (WT or mutant) constructs or co-transfected with TRPV4-GFP and RhoA-Myc (WT or mutant) constructs using Lipofectamine LTX. Calcium imaging was performed on a Zeiss Axio Observer.Z1 inverted microscope equipped with a Lambda DG-4 (Sutter Instrument Company, Novato, CA) wavelength switcher. Cells were bath-loaded with Fura-2 AM (8 μM, Life Technologies) for 45-60 min at 37°C in calcium-imaging buffer (150 mM NaCl, 5 mM KCl, 1 mM MgCl2, 2 mM CaCl2, 10 mM glucose, 10 mM HEPES, pH 7.4). For hypotonic saline treatment, one volumes of NaCl-free calcium-imaging buffer was added to one volume of standard calcium-imaging buffer for a final NaCl concentration of 70 mM. Cells were imaged every 10 sec for 30 sec prior to stimulation with hypotonic saline or GSK101, and then imaged every 10 sec for an additional 2 min. Calcium levels at each time point were computed by determining the ratio of Fura-2 AM emission at 340 nM divided by the emission at 380 nM. Data were expressed as Fura ratio over time.

### Data availability

Coordinates have been deposited in the Protein Data Bank with the PDB IDs - 8FC9 (TRPV4-RhoA, apo), 8FC7 (TRPV4-RhoA, GSK279-bound closed), 8FCB (TRPV4-RhoA, GSK101 bound, open), 8FC8 (TRPV4, GSK101 bound, open), 8FCA (TRPV4, 4α−PDD-bound, putative open) respectively. The cryo-EM maps have been deposited in the Electron Microscopy Data Bank with the IDs EMD - 28977 (TRPV4-RhoA, apo), 28975 (TRPV4-RhoA, GSK279-bound closed), 28976 (TRPV4-RhoA and TRPV4 only, GSK101 bound, open), 28978 (TRPV4, 4α−PDD-bound, putative open), respectively.

## Acknowledgements

Cryo-EM data were screened and collected at the Duke University Shared Materials Instrumentation Facility (SMIF) and at the Pacific Northwest Center for Cryo-EM (PNCC) at OHSU. We thank Nilakshee Bhattacharya at SMIF and Janette Myers at PNCC for assistance with the microscope operation. We thank Ying Yin and Justin Fedor for critical manuscript reading and advice throughout the project and Yang Suo for assistance in data collection. This research was supported by National Institutes of Health grants R35NS097241 (S.-Y.L.), R35NS122306 (C.J.S) and K08NS102509 (B.A.M.) and Muscular Dystrophy Association 629305 (C.J.S.). A portion of this research was supported by NIH grant U24GM129547 and performed at the PNCC at OHSU and accessed through EMSL (grid.436923.9), a DOE Office of Science User Facility sponsored by the Office of Biological and Environmental Research. DUKE SMIF is affiliated with the North Carolina Research Triangle Nanotechnology Network, which is in part supported by the NSF (ECCS-2025064).

**Author Contributions**: D.K. conducted biochemical preparation, sample freezing, cryo-EM data collection, and processing, F.Z. conducted biochemical preparation and electrophysiology experiments, all under the guidance of S.-Y.L. D.K. and S.-Y.L. performed model building and refinement. B.A.M, and M.K. carried out immunoprecipitation and calcium imaging experiments under the guidance of C.J.S. S.-Y.L. D.K. C.J.S. B.A.M. J.M.S. and F.Z. wrote the paper.

**Competing Interests**: The authors declare no competing interests.

---

**Extended Data Fig. 1.**
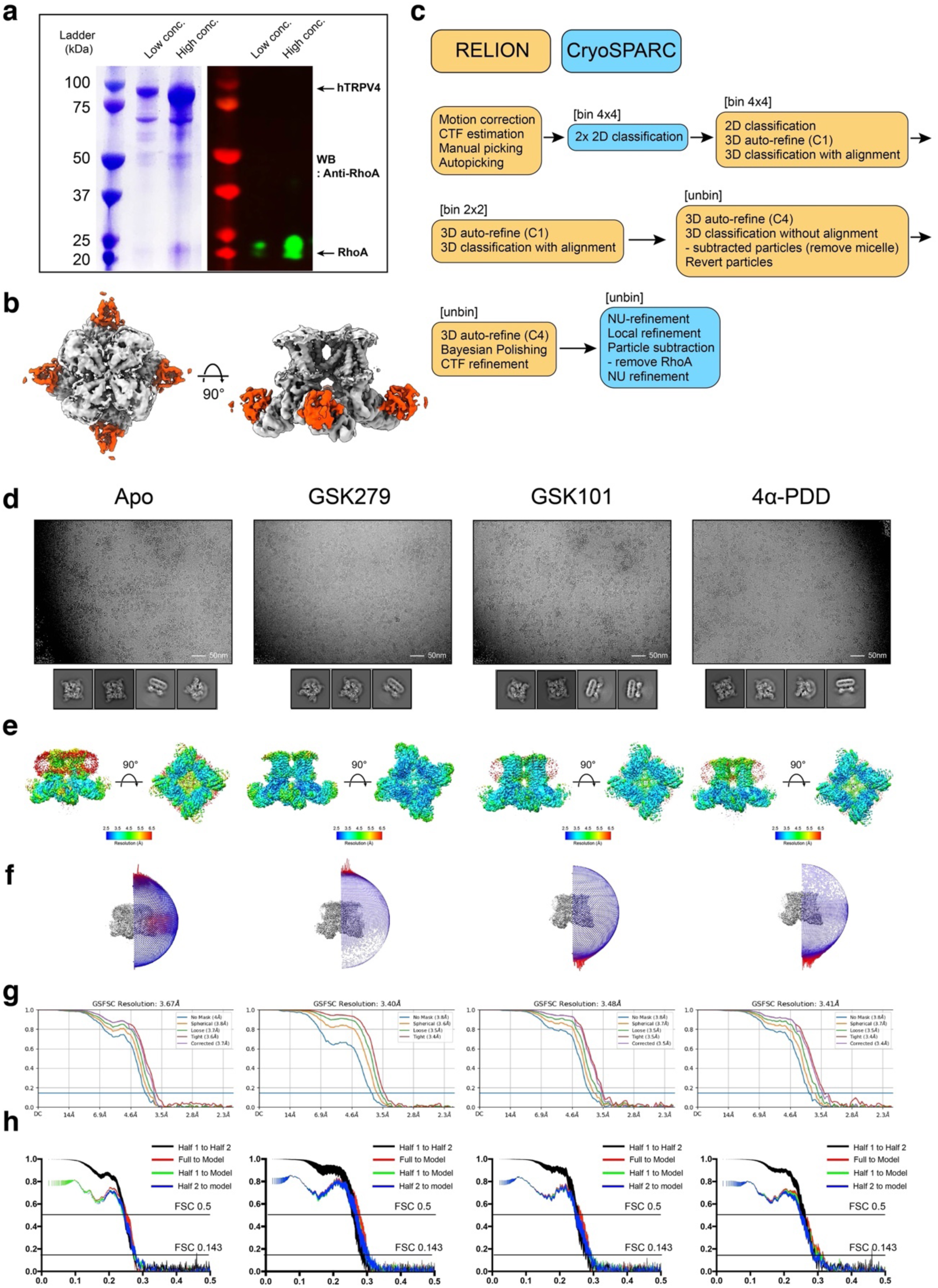
Cryo-EM data processing summary. **a**, SDS-PAGE and western blot analysis of purified hTRPV4-RhoA. **b**, Cryo-EM reconstruction of GSK279-hTRPV4-RhoA imposing symmetry (C1 symmetry). The red-colored densities indicate RhoA binds to each protomer. **c**, General cryo-EM data processing procedure. Tasks performed in the RELION software are colored in orange and those in the cryoSPARC in blue. **d**, Representative micrographs of hTRPV4-RhoA sample in vitreous ice and 2D classification images. **e**, Local resolution estimations. **f**, Euler distribution plots. **g**, Fourier shell correlation (FSC) curves of the final 3D reconstructions with different masking, as calculated in cryoSPARC. **h**, Map-to-model correlation plots for both full maps and half maps.

**Extended Data Fig. 2.**
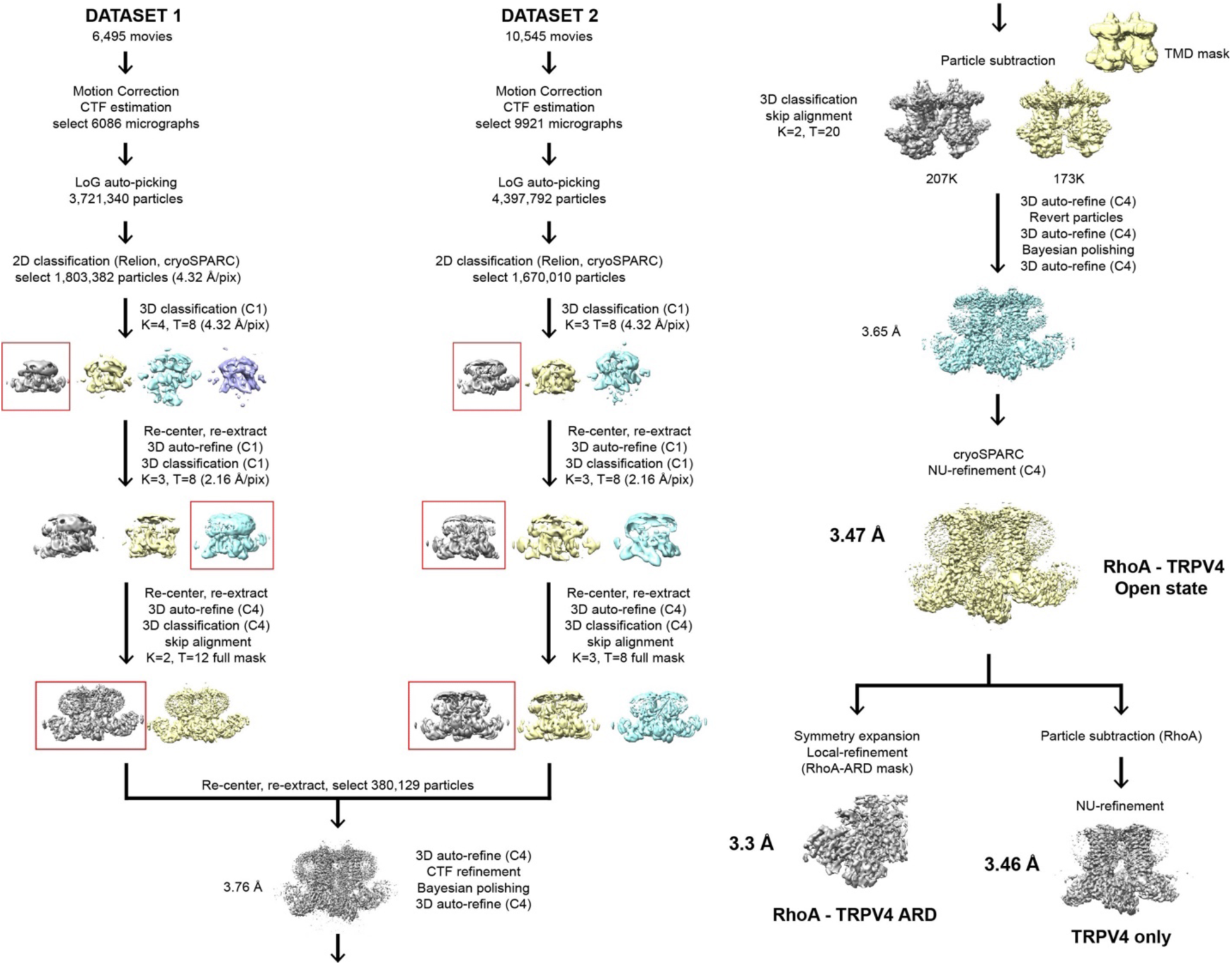
Cryo-EM data processing of the open state (GSK101) structure. The detailed procedure for determining the open state GSK101-hTRPV4-RhoA structure. The final cryo-EM reconstruction at a resolution of 3.47 Å of GSK101-hTRPV4-RhoA, RhoA-ARD focused map at a 3.30 Å, TRPV4 focused map at a 3.46 Å.

**Extended Data Fig. 3.**
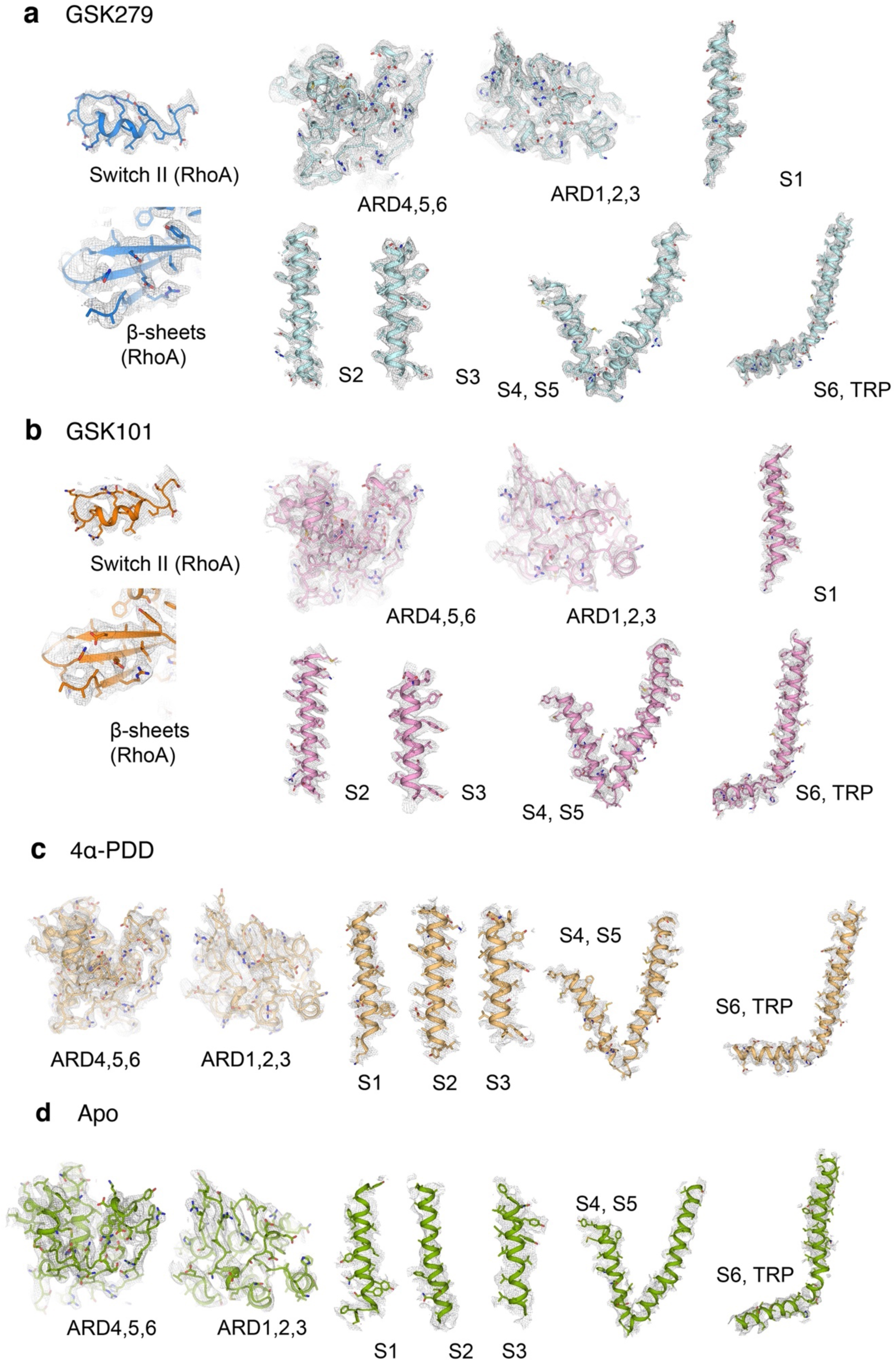
Representative Cryo-EM density of the TRPV4-RhoA structures. **a-d**, Representative cryo-EM densities for various structural elements in the structures of GSK279-hTRPV4-RhoA (**a**, blue and cyan), GSK101-hTRPV4-RhoA (**b**, orange and pink), 4α-PDD-hTRPV4-RhoA, (**c**, light gold), and apo-hTRPV4-RhoA (**d**, green). RhoA densities in the GSK279- and GSK101-bound structures are from the focused-refined maps. EM densities are shown as gray meshes at thresholding of 0.23-0.25, 0.22–0.24, 0.17–0.19, and 0.16–0.18, respectively, in **a-d**.

**Extended Data Fig. 4.**
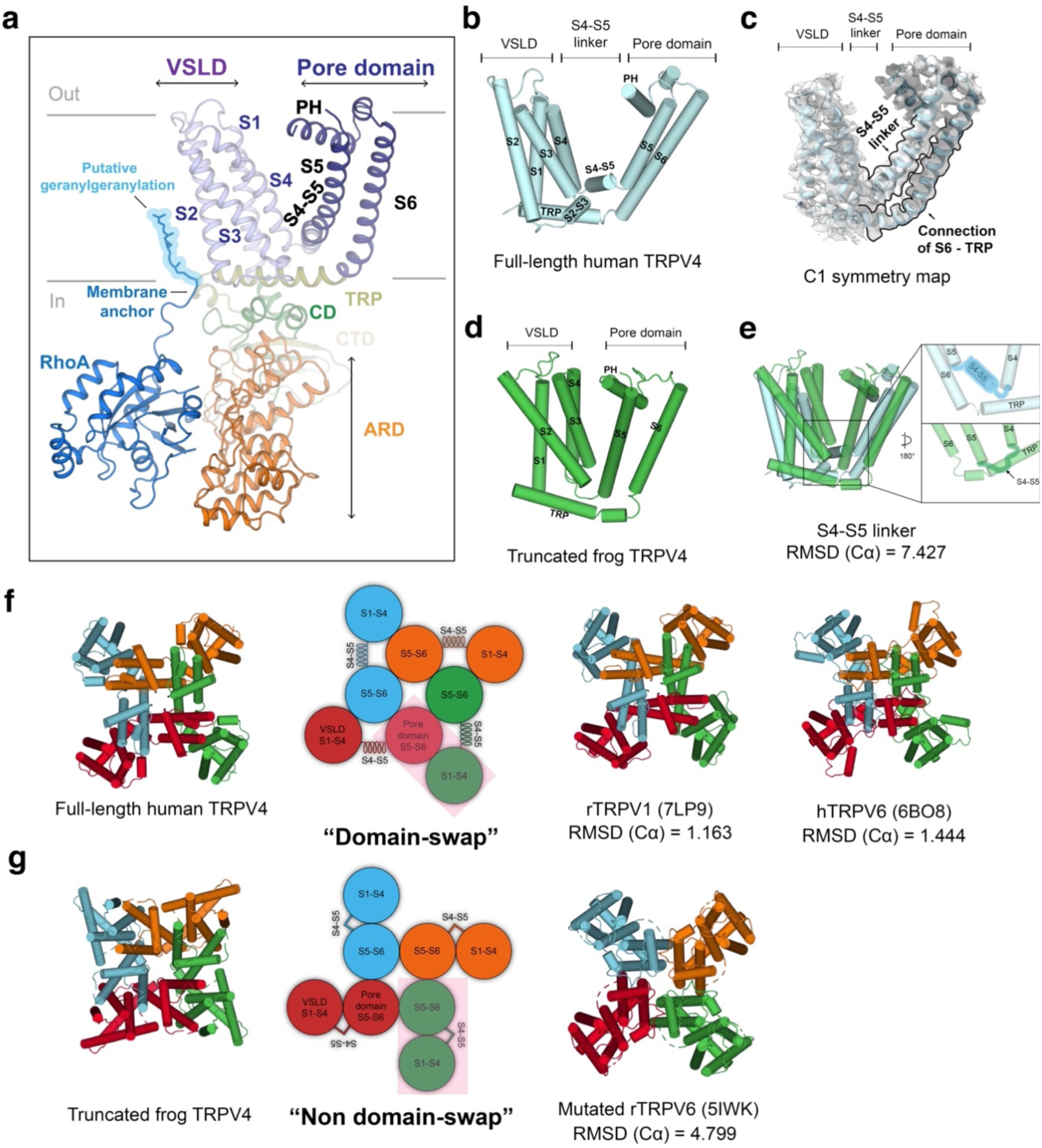
Structural comparisons between human TRPV4 and frog TRPV4. **a**, Architecture of the hTRPV4 protomer and RhoA with subdomains indicated: ankyrin repeat domain (ARD), coupling domain (CD), transmembrane helices (S1-S6), pore helix (PH), TRP helix, C-terminal domain (CTD), and RhoA. Membrane-anchored lipid tail of RhoA is highlighted in blue. **b**, Cylinder representation of the TMD of human TRPV4 in cyan. **c**, Cryo-EM density (no imposed symmetry) of GSK279-hTRPV4-RhoA complex for the domain-swapping linkers (S4-S5 and S6-TRP), at thresholding 0.35. **d**, Cylinder representation of the TMD of truncated frog TRPV4 in green. **e**, Comparison of the TMDs of human TRPV4 and frog TRPV4 viewed from the membrane plane. The linkers between S4 and S5 are highlighted. **f**,**g**, Comparison of domain-swapping architectures of human TRPV4, rat TRPV1, human TRPV6 (**f**), truncated frog TRPV4, and mutated rat TRPV6 (**g**).

**Extended Data Fig. 5.**
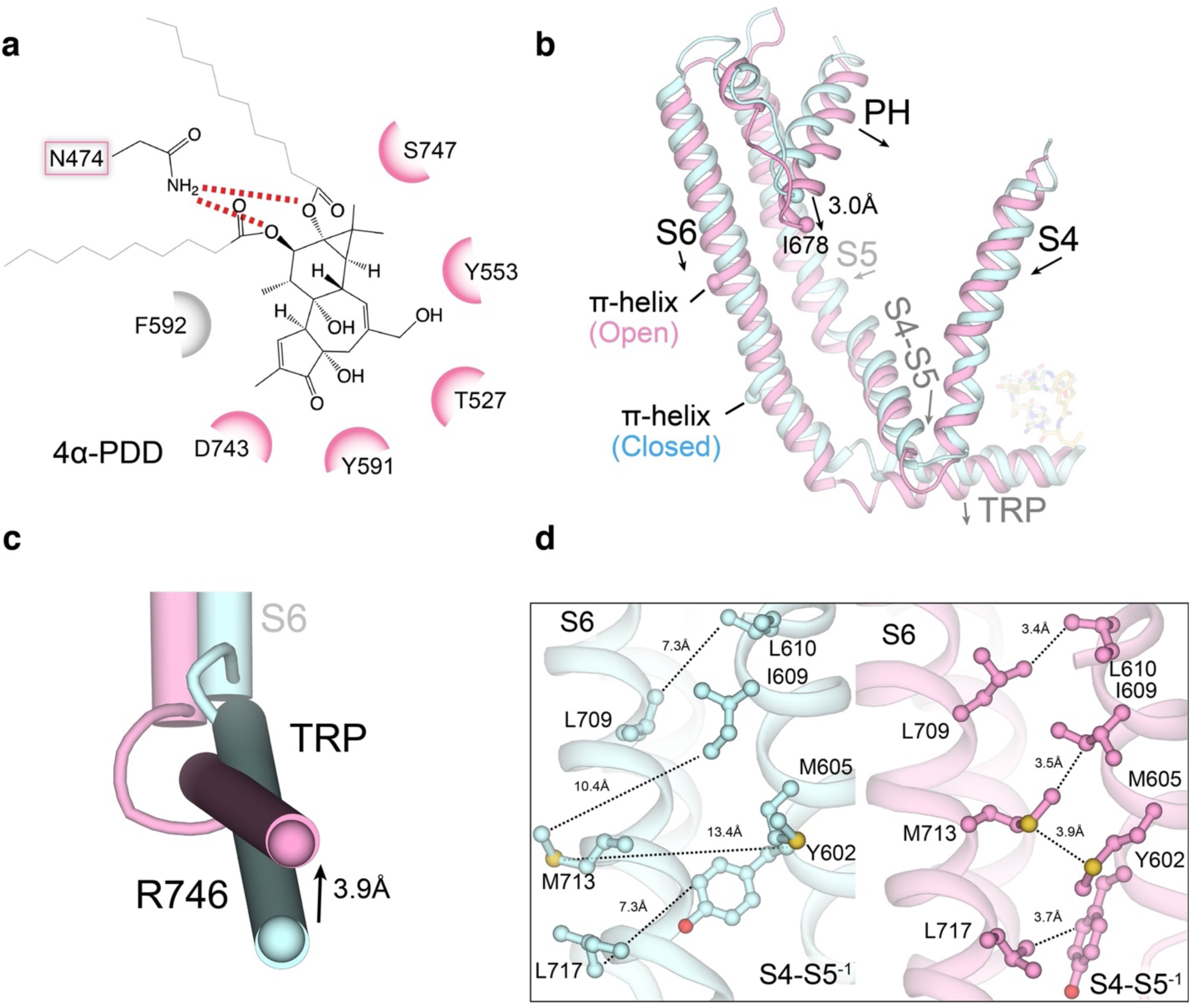
Structural comparison of hTRPV4-RhoA inter- and intra-subunit interfaces in the open and closed states. **a**, Ligplot^62^ schematics of the 4α-PDD interaction with hTRPV4, with key chemical positions labelled. Pink colored residues interact with both GSK101 and GSK279. **b**, Comparison of conformational changes at pore-domain of GSK279-hTRPV4-RhoA (cyan) and GSK101-hTRPV4-RhoA (pink). Ligands are shown in sticks. Arrows indicate helix movement with distances indicated between reference points (as spheres). **c**, Closed-up view of S6b and the TRP helix. Arrows indicate helical movements with distances indicated between reference points (as spheres). **d**, Side-by-side comparison of the closed (cyan) and open states (pink) at inter-subunit interfaces. Sidechains are shown as sticks. Dashed lines indicate the distances between corresponding residues.

**Extended Data Fig. 6.**
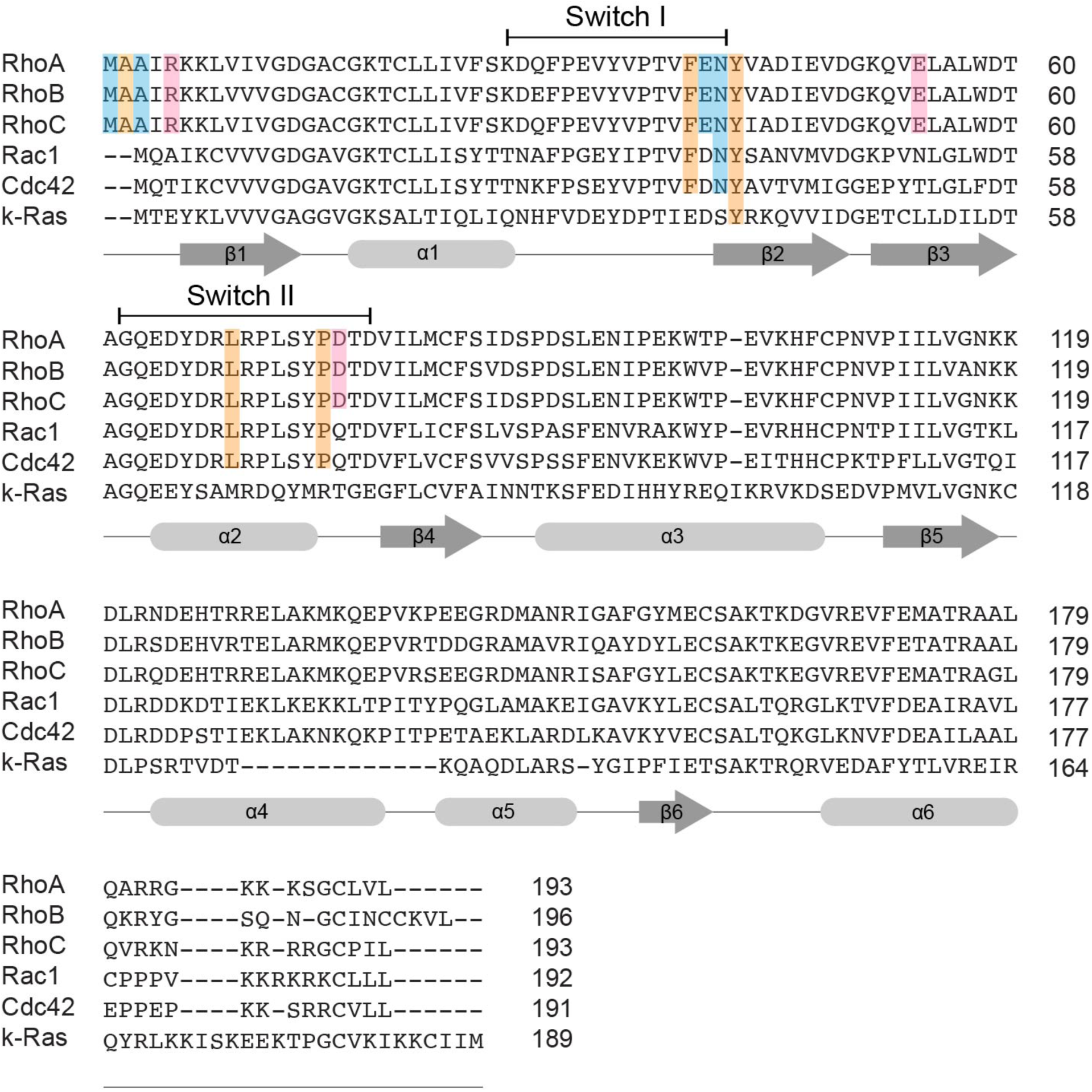
Sequence-alignment of Rho family of GTPases. Secondary structure elements based on RhoA are shown as gray cylinders (helices) and arrows (beta-strand). The RhoA residues critical for interaction are highlighted: blue for backbone - M1, A3, E40, N41, pink for sidechain - R5, E54, D76, and orange for hydrophobic - A2, F39, Y42, L69, P75.

**Extended Data Fig. 7.**
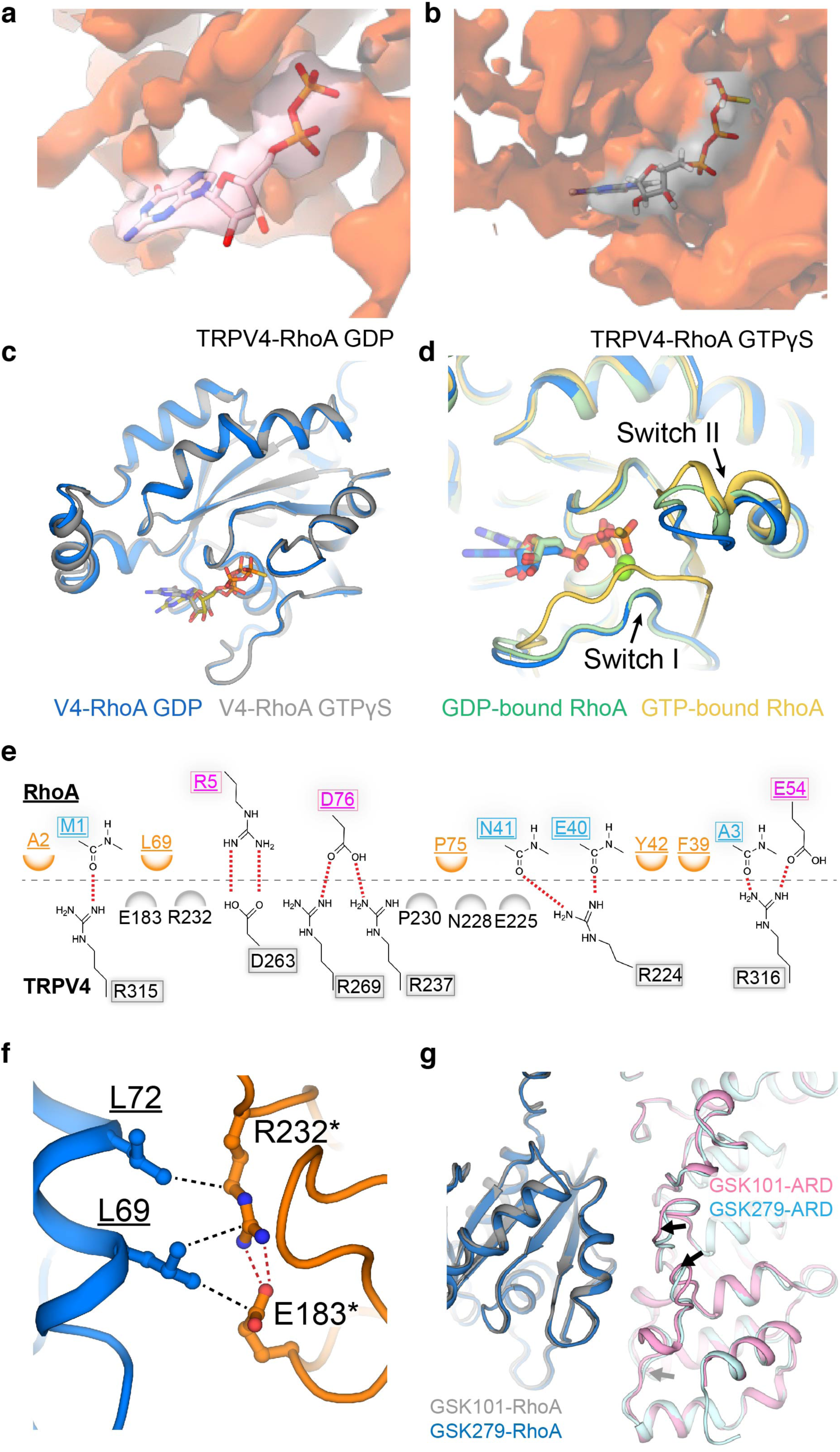
Structural comparisons of the TRPV4-bound RhoA and published RhoA. **a**,**b**, Close-up view at the nucleotide binding site in RhoA of 3D reconstructions of GSK279-hTRPV4-RhoA-GDP (**a**) and GSK101-hTRPV4-RhoA-GTPγS (**b**) at thresholding 0.17 and 0.135, respectively. Nucleotides are shown as sticks. **c**, Comparison of RhoA conformations of GSK279-hTRPV4-RhoA-GDP and GSK101-hTRPV4-RhoA-GTPγS. Nucleotides are shown as sticks. **d**, Comparison of RhoA conformations of GSK279-hTRPV4-RhoA-GDP (blue), GDP-bound RhoA (PDB-1FTN, green), and GTPγS-bound RhoA (PDB-1A2B, gold). Nucleotides are shown as sticks and Magnesium ions are shown as spheres. **e**, DIMplot^62^ schematics of TRPV4 ARD and RhoA interactions, with critical positions labeled. Pink-colored residues indicate interactions via sidechains. Blue-colored residues indicate backbone-sidechain interactions. Orange-colored residues indicate hydrophobic interactions. **f**, Detailed hydrophobic interaction of TRPV4-ARD and RhoA. Underlined residues located in RhoA, and annotated residues indicate disease-causing mutations. The red and black dotted lines indicate salt bridges and hydrophobic interactions, respectively. **g**, Comparison of the interaction interface between GSK279-hTRPV4-RhoA-GDP and GSK101-hTRPV4-RhoA-GTPγS. Arrows indicate conformational changes.

**Extended Data Fig. 8.**
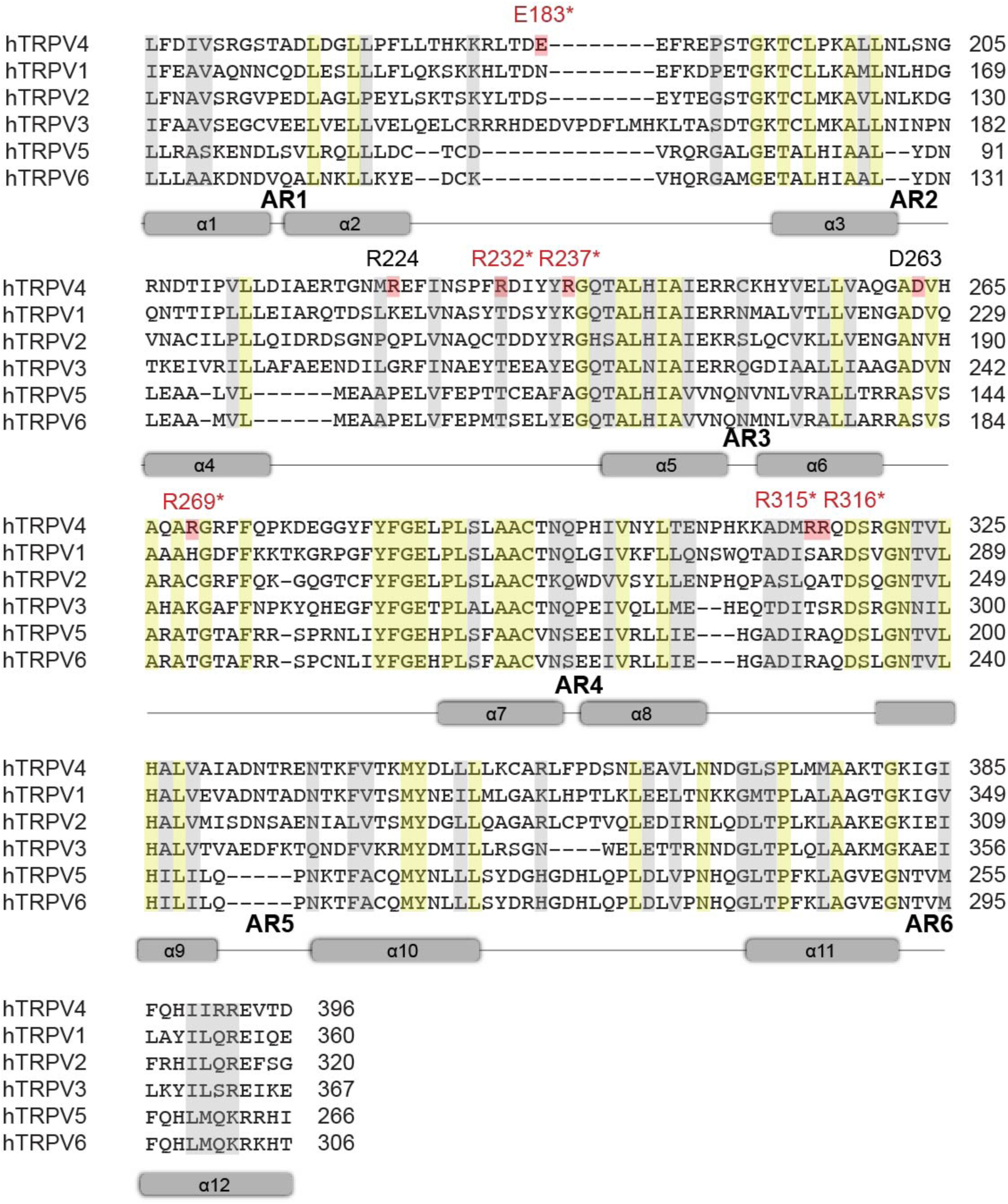
Sequence alignment of the ARDs in human TRPV channels. The α-helices shown as gray cylinders, well-conserved residues highlighted in yellow (identical) and gray (similar), and key residues for TRPV4 interaction are colored in red. Annotated residues are variant positions in neuropathy disease.

**Extended Data Fig. 9.**
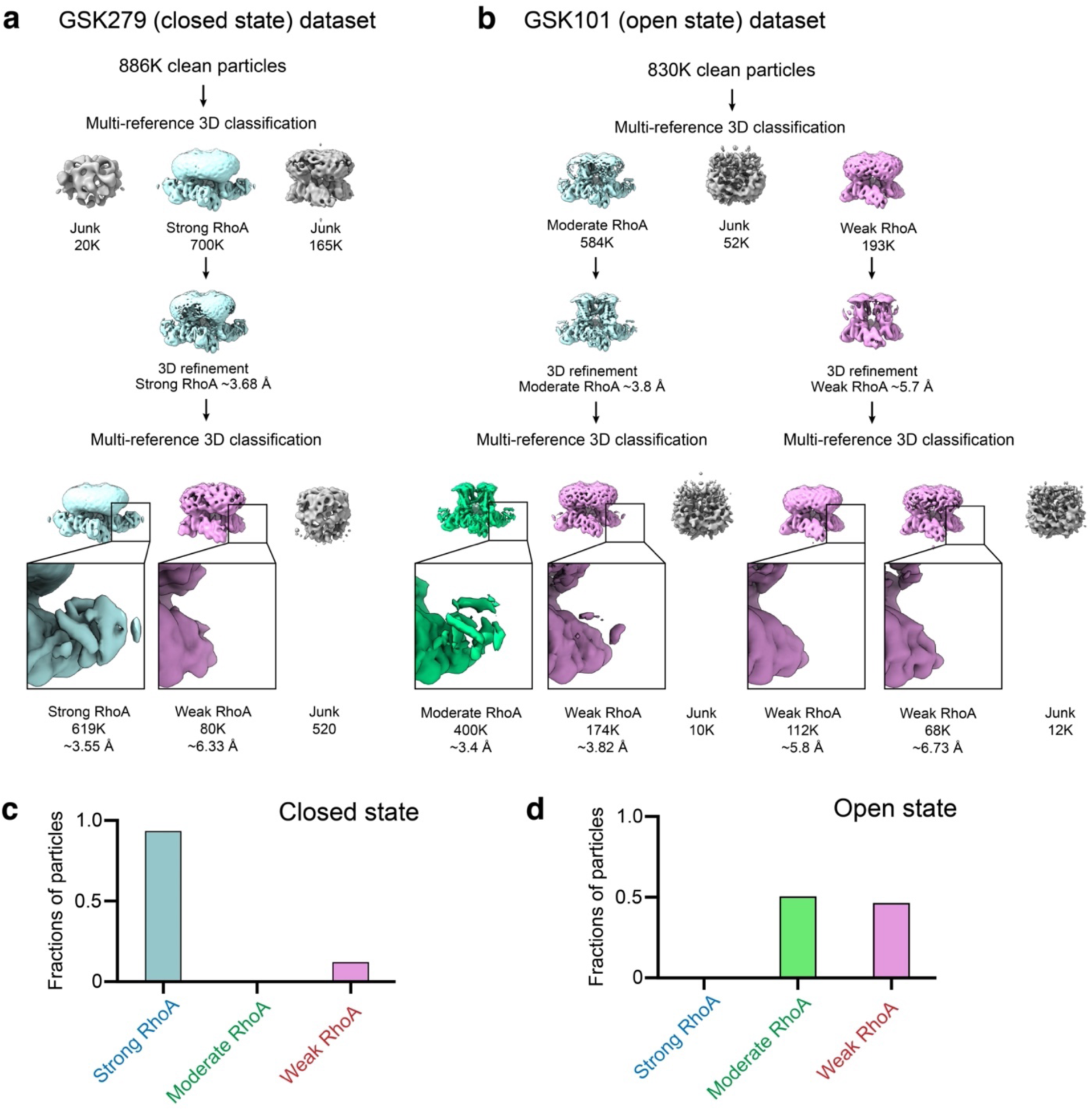
Image processing workflow for the RhoA-bound fraction analysis. **a**, Data processing workflow of GSK279-hTRPV4-RhoA dataset for the purpose of the RhoA-bound fraction analysis. 3D reconstruction thresholding of 0.58. **b**, Data processing workflow of GSK101-hTRPV4-RhoA dataset for the purpose of the RhoA-bound fraction analysis. The red- and blue-outlined subsets indicate the RhoA-bound fraction and RhoA-unbound fraction, respectively. 3D reconstruction thresholding of 0.58. **c**, Class distributions of particles for GSK279-hTRPV4-RhoA. **d**, Class distributions of particles for GSK101-hTRPV4-RhoA.

